# Future Preventive Gene Therapy of Polygenic Diseases from a Population Genetics Perspective

**DOI:** 10.1101/770396

**Authors:** Roman Teo Oliynyk

**Affiliations:** Centre for Computational Evolution, University of Auckland, Auckland 1010, New Zealand; Department of Computer Science, University of Auckland, Auckland 1010, New Zealand

**Keywords:** polygenic risk, PRS, CRISPR, heritability, polygenic disease, simulation, gene therapy, stratification, lifetime risk, admixture

## Abstract

With the accumulation of scientific knowledge of the genetic causes of common diseases and continuous advancement of gene-editing technologies, gene therapies to prevent polygenic diseases may soon become possible. This study endeavored to assess population genetics consequences of such therapies. Computer simulations were used to evaluate the heterogeneity in causal alleles for polygenic diseases that could exist among geographically distinct populations. The results show that although heterogeneity would not be easily detectable by epidemiological studies following population admixture, even significant heterogeneity would not impede the outcomes of preventive gene therapies. Preventive gene therapies designed to correct causal alleles to a naturally-occurring neutral state of nucleotides would lower the prevalence of polygenic early- to middle-age-onset diseases in proportion to the decreased population relative risk attributable to the edited alleles. The outcome would manifest differently for late-onset diseases, for which the therapies would result in a delayed disease onset and decreased lifetime risk, however the lifetime risk would increase again with prolonging population life expectancy, which is a likely consequence of such therapies. If gene therapies that prevent heritable diseases were to be applied on a large scale, the decreasing frequency of risk alleles in populations would reduce the disease risk or delay the age of onset, even with a fraction of the population receiving such therapies. With ongoing population admixture, all groups would benefit over generations.

## 1. Introduction

Research into the causality and liability of diseases primarily based on familial and populational observations greatly pre-dates the discovery of DNA structure and the genetic code in 1953 by Watson and Crick [1]. Initially, it was only possible to estimate the frequency of highly malignant mutations in human populations [2]. It took several decades for experimental techniques to develop sufficiently to sequence the human genome [3]. Whole genome sequencing (WGS) and genome-wide association studies (GWASs) have provided experimental insights into the genetic architecture of polygenic diseases that could be only hypothesized a decade or two earlier [4].

The search for singular genetic mutations started decades ago and continued with GWASs and WGS, which led to the discovery of many thousands of highly malignant so-called Mendelian conditions. Among such conditions are sickle-cell anemia, Tay–Sachs disease, cystic fibrosis, hemophilia, thalassemia, Huntington disease, early-onset Alzheimer’s disease, and macular degeneration, as well as mutations in the *BRCA*1/2 genes, which are causally linked to multiple types of cancer, especially breast cancer [5]. On its own, the prevalence of each such disease in the population is relatively low. The mutations that cause the majority of Mendelian conditions are known and usually involve single nucleotide variants (SNVs) that are associated with a high susceptibility to these diseases, with other sequence rearrangements representing an aggregate 13% of mutations [6,7]. The OMIM Gene Map Statistics [5] database lists over 4000 of such gene mutations responsible for almost 6500 phenotypic conditions or syndromes, and The Human Genome Mutations Database [8] lists more than 250,000 disease-causing mutations. It has been estimated that, on average, an individual carries 0.58 recessive alleles that can lead to complete sterility or death by reproductive age when homozygous [9]. The fact that this number is an average of a large variety of very rare mutations distributed throughout the genome indicates that severe events, which occur when these rare alleles affect a particular gene pair in one descendant, are an infrequent occurrence. However, in aggregate, less malignant diseases caused by rare mutations affect a noticeable fraction of the population, with approximately 8% of individuals affected [7,10].

Tests have been conducted on many experimental gene therapy techniques that target diseases typically caused by a single defective gene or SNV. Ginn *et al*. [11] identified 287 trials that had been performed by the end of 2017 on inherited monogenic disorders, with the overall number of clinical trials of gene therapies, predominantly in the oncology field, exceeding 2600. Philippidis [12] summarized 25 gene-editing therapies that were under clinical trial during the first quarter of 2019. All therapies in these studies focused exclusively on the clinical or reactive—rather than prophylactic—treatment of genetic conditions. Although not yet technologically or medically possible, the potential of applying germline gene-editing therapy to prevent at least some of these diseases is being increasingly discussed. Public understanding of the expected health benefits of such therapies is gradually building [13,14], and is notably present in the recommendations of the UK Nuffield Council on Bioethics [15] report *Genome editing and human reproduction: Social and ethical issues* (2018). Hypothetically, when the medical technology becomes available to safely and accessibly correct these mutations, and if governmental regulations allow it in the future [16], treated individuals and their descendants (in cases of heritable gene therapies) will be effectively cured and have no need for concern about the single specific cause of their disease.

In contrast to Mendelian conditions, polygenic or complex disease liability is attributed to hundreds and thousands of gene variants or single nucleotide polymorphisms (SNPs) of typically small effect that, in combination, constitute the polygenic disease risk of an individual [17–19]. The polygenic risk score (PRS) of an individual at higher risk for a polygenic disease reflects the presence of a higher number of detrimental gene variants [20] relative to the average distribution of common gene variants in the population. Polygenic diseases include highly prevalent old-age diseases—termed late-onset diseases (LODs)—that eventually affect most individuals (for example, cardiovascular disease, particularly coronary artery disease, cerebral stroke, type 2 diabetes, senile dementia, Alzheimer’s disease, cancers, and osteoarthritis) [21–28], as well as earlier-onset diseases and phenotypic features such as susceptibility to asthma and psychiatric disorders and particular height and high body mass index (BMI) characteristics [4]. Over the past ten years, GWAS results have been reported for hundreds of complex traits across a wide range of phenotypes. These studies have led to a well-established consensus that a large number of common low-effect variants can explain the heritability of the majority of complex traits and diseases [4,29,30]. Post-GWAS epidemiological studies of gene–environment interactions have generally reported multiplicative joint associations between low-penetrant SNPs and environmental risk factors, with only a few exceptions [31].

Geographic and local population genetic stratification and variation complicate the ability to diagnose and treat medical conditions [32] (for additional exposition, see Addendum A.1). The predictive utility of GWAS and GWAS PRSs also varies broadly if the risk score is applied to a population other than the one for which the score was initially determined [33–35]. At the same time, there are many indications of the commonality of causal gene variants for polygenic diseases among geographically distinct populations [36,37], while admixed populations present an intermediate liability to diseases [38–40].

Even when the majority of causal gene variants are common among populations, they are difficult to match precisely in genetically stratified populations for two main reasons. First, the GWAS PRS is composed of representative so-called “tag” SNPs. Rather than being true causal variants, tag SNPs are from a genomic region that exerts a single or combined effect of multiple detrimental and protective SNPs in various degrees of linkage disequilibrium and varying allele frequencies in different subpopulations [41,42] (see Figure 1A). Thus, although only a small fraction of true causal SNPs for each polygenic condition have been identified, PRSs can be determined since they rely on an aggregate of implicit determinations that are likely to significantly differ among the population-specific background of non-causal SNPs [41]. The second reason that underlies this challenge is that, in addition to differences in SNPs, there are less-researched structural variations that differ among populations and can influence disease liability [43]. Major projects are underway that aim to comprehensively catalog the detrimental structural variation in diverse populations [44]. In parallel, the advancement of biomedical techniques will facilitate the detection of germline structural variants for clinical validation and research in the future [45].

**Figure 1.**
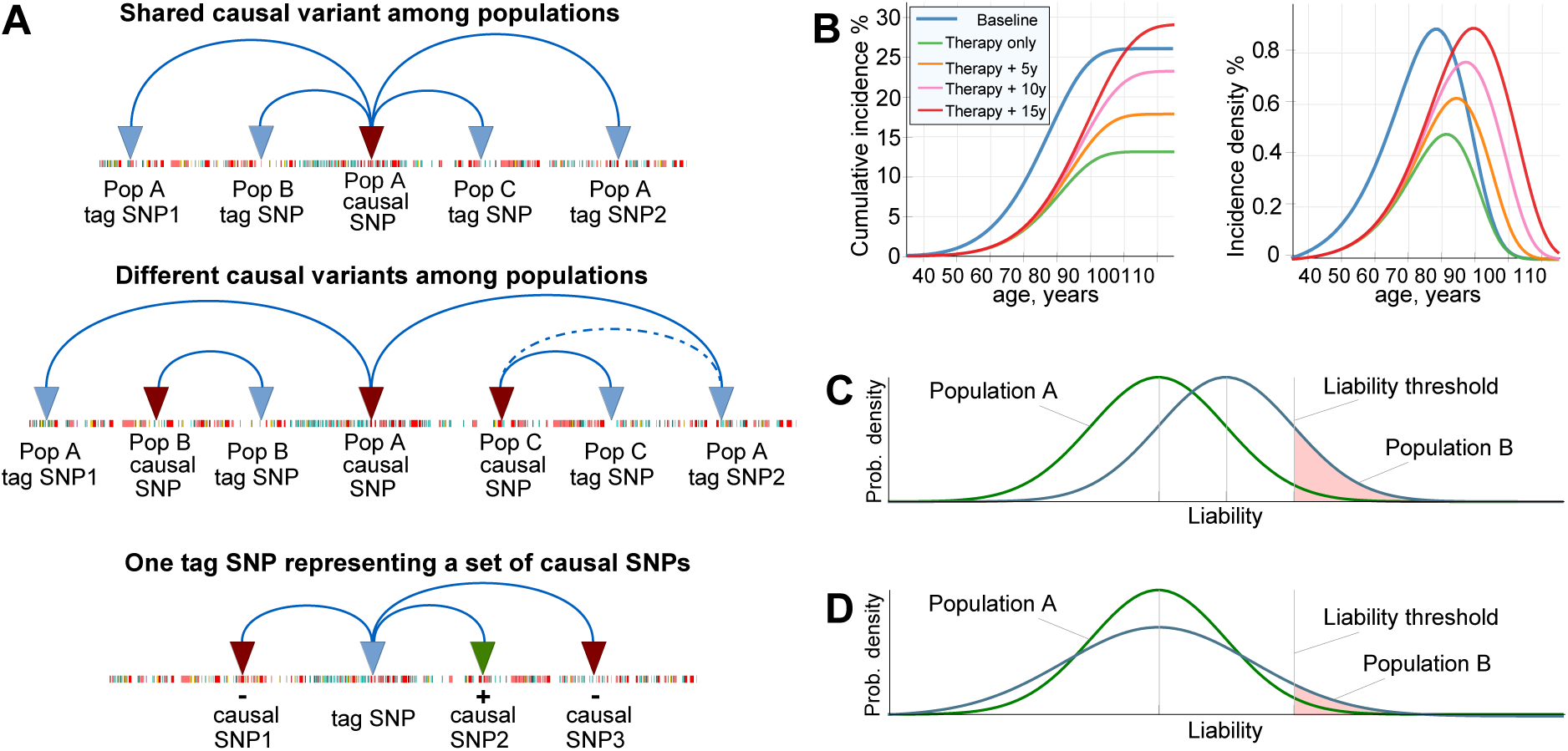
Illustrations to the concepts in this Introduction. (A) Genome-wide association study (GWAS) assignment of tag single nucleotide polymorphisms (SNPs) differs in geographically diverse populations because of differences in linkage disequilibrium and study setup. The same causal SNPs can be assigned to different tag SNPs in different populations (and subpopulations), and different causal SNPs may overlap in varied ways among populations. Tag SNPs can represent a set of causal SNPs that are protective (+), detrimental (-), or both [34]. (B) Lifetime risk and incidence density distribution of late-onset disease (LOD) under Cox’s proportional hazards/multiplicative polygenic risk model. The example shown is coronary artery disease, where the LOD lifetime risk is delayed after therapy that lowers the population polygenic risk, and the lifetime risk is regained with increasing life expectancy [46]. (C) Falconer’s liability threshold model with different mean liabilities and the same variance (*Prob. density* stands for the probability density of an individual succumbing to a disease) [47,48]. Under this model, the disease prevalence is a function of the disease liability (as termed by Falconer), which can be understood as the polygenic risk score of true causal gene variants. For Population B, the area to the right of the liability threshold is larger, as is the disease prevalence; the vertical liability threshold line is the initial Falconer interpretation for illustration purposes. Modern approaches can be perused in [49]. (D) Falconer’s liability threshold model with the same mean liability and different liability variances. If both distributions are normalized, the prevalence will be larger for a wider variance, particularly distinct for smallest prevalence values, and it will remain identical between populations A and B at a prevalence of 50%.

For LODs, a combination of genetic liability, environmental factors, and the physiological decline of multiple organ systems leads to individual disease presentations [27]. Earlier research evaluated the risk allele distributions that accompany aging for polygenic LODs [50] to quantify the potential of future preventive gene therapies to delay the onset age and lifetime risk of such LODs [46]. This is demonstrated in Figure 1B, leveraging age-specific incidence rates under Cox’s proportional hazards model [31,51]. A recent clinical data analysis confirmed these theoretical predictions [52].

The polygenic diseases with highest incidence in early- and middle-age, that are the focus of the current research, are exemplified by asthma [53,54], chronic migraine [55,56], Dupuytren’s disease [57], rheumatoid arthritis [58], lupus erythematosus [59], schizophrenia and bipolar disorder [60], and Crohn’s disease [61,62]. The lower prevalence of these diseases contrasts with the high prevalence of some LODs, highlighting differences in their evolutionary and causal manifestations [63]. These diseases are less suitable for the age-specific rates approach [46], because subjects with an earlier age at disease onset do not necessarily show an increased polygenic risk burden, as exemplified by schizophrenia incidence [64]. The liability to these diseases is often illustrated using the liability threshold model proposed by Falconer [48] (see Figure 1(C–D)).

In this study, computer simulations were used to evaluate the magnitude of the heterogeneity in alleles causal for polygenic diseases that could exist among geographically distinct populations. The results show that even with significant heterogeneity, the outcomes of preventive gene therapies would not be impeded. Population genetics simulations were performed for representative scenarios of preventive gene therapies designed to turn true causal alleles into a naturally existing neutral state of nucleotides for polygenic **E**arly- to **M**iddle-age-**O**nset **D**iseases (EMODs). The simulations determined that the disease prevalences would decrease proportionately to decrease in the average population relative risk attributable to the edited alleles, and evaluated the progression of population admixture that would accompany such therapies. The combination of these EMOD findings with earlier published LOD conclusions resulted in a comprehensive picture of preventive polygenic disease gene therapy from a population genetics perspective.

## 2. Results

### 2.1 Admixture of Populations with Matching Mean PRSs: To What Extent Can Causal Risk Alleles of Polygenic Diseases Differ Between Populations?

The first set of simulations evaluated the blending admixture of two simulated populations with equal liability to a disease. The disease heritability was set at 50%, the mid-range heritability of polygenic diseases [65,66]. The disease SNP sets were built using the common low-effect genetic architecture, and the population genetics simulation progressed through generations. Four simulated scenarios, in which the combined effect of SNPs differed between the populations by 100%, 65%, 33%, and 20%, were considered.

The simulations recorded the changes in the variance of the population PRS and disease prevalence as generations progressed. The simulated diseases were polygenic EMODs, which are model polygenic diseases whose maximum incidence occurs at young- to middle-age, with a negligible incidence at older ages. In this publication, the term “prevalence”, used in reference to EMODs, always means the prevalence at an age later than the typical age of onset range.

The results presented in Figure 2 show that for all scenarios of differing SNP architectures, the PRS variance gradually increased starting from the second admixed generation, and it continued to increase in subsequent generations. The consequences of this pattern are illustrated in Figure 1D. The rise was gradual, resulting in the fifth generation in a 3% increase in prevalence for the scenario in which all causal SNPs differed between the populations, and it increased by just a fraction of a percent for the scenario in which one-fifth of causal SNPs differed. By the 25th generation, the prevalence values for the highest and lowest differences in genetic architecture causality scenarios were 1.12% and 1.03%—a 12% with a 3% increase in the prevalence in the populations before admixture. These results are summarized in Table 1. The gradual increases in variance and prevalence were due to gradual recombination of the population genome. Figure 2D shows the result of accelerating the recombination to 1000 crossovers per genome per generation. In this figure, the population risk variance and prevalence approach the equilibrium within a few generations. This increase in variance with the admixture of diverse populations was previously reported with much smaller magnitudes of causal allele stratification based on actual allele frequencies in human populations [67].

**Figure 2.**
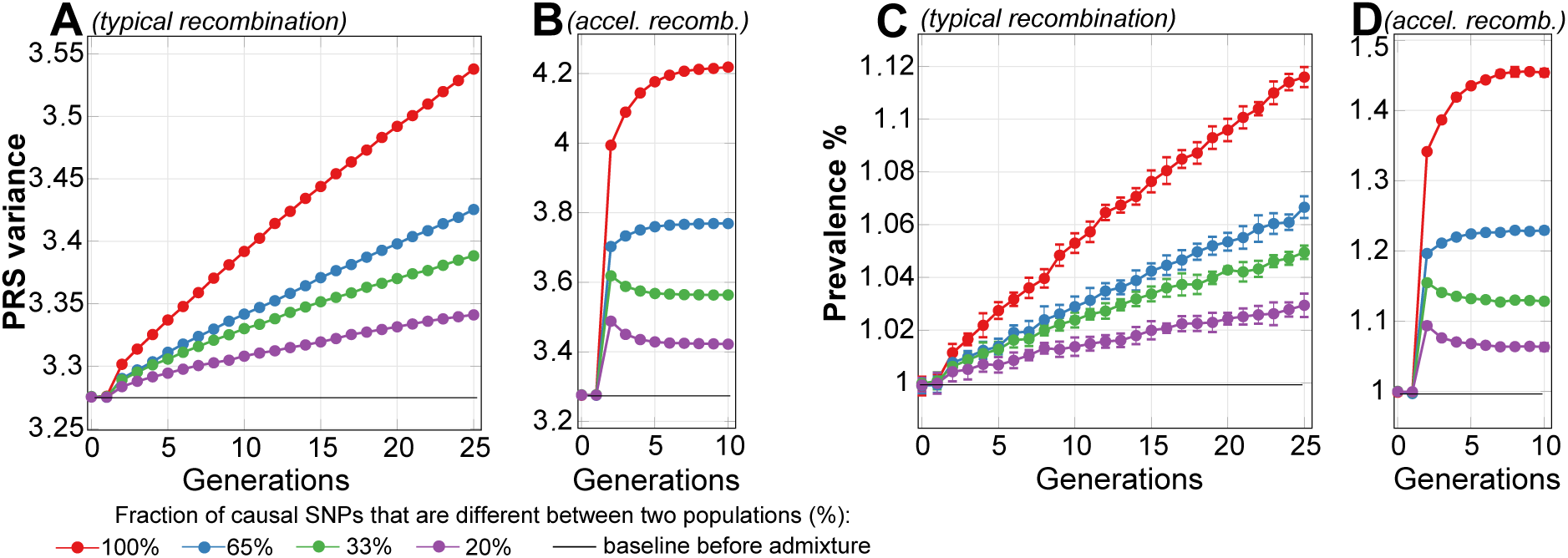
The admixture of two simulated populations with equal liability to a disease with 50% heritability and 1% prevalence. The plots represent four scenarios, in which causal SNPs differ between the populations in a range between 100% (all causal SNPs for this disease are different between the two populations) and 20% different (one-fifth of causal SNPs are different, with the remaining majority of causal SNPs in common), as listed in the figure legend. The blending commences at generation 2. (**A**) represents the change in variance of the polygenic risk score (PRS) as a result of 100% blending of two equally sized populations over 25 generations with a relatively typical recombination rate of 36 recombinations per parental genome; (**B**) shows accelerated recombination (accel.recomb.) in which 1000 recombinations were applied per parental genome, resulting in the variance level quickly stabilizing to equilibrium; (**C**) represents the change in the prevalence of the disease with a baseline prevalence of 1%, corresponding to the variance change in the previous plot; (**D**) shows accelerated recombination (accel.recomb.) in which 1000 recombinations were applied per parental genome, resulting in the prevalence level quickly stabilizing to equilibrium.

**Table 1.** Summary of the admixture of two simulated populations with equal liability to a disease and varied fractions of differing causal SNPs.

| Fraction of differing causal SNPs | 100% | 65% | 33% | 20% |
| --- | --- | --- | --- | --- |
| Second-generation prevalence increase, % | 1 | <1 | <1 | <1 |
| Fifth-generation prevalence increase, % | 2.7 | 1.4 | 1.3 | 0.8 |
| Asymptotic prevalence increase limit, % | 45 | 22 | 11 | 6.3 |

The simulated values are for a disease with early- to middle-age onset, 50% heritability, and a 1% prevalence/lifetime risk. The relative prevalence increase is calculated in comparison to the baseline prevalence, where, for example, the prevalence increase from 1% to 1.45% represents a 45% relative increase.

This phenomenon can be simplistically explained using an example of two risk alleles, each unique to one of two identically sized populations with identical disease risk. When these populations blend together, the frequency of risk alleles is expected to be average in the resulting population, with the resulting average effect size, or PRS, remaining unchanged. At the same time, following Equation (A1), the sum of the variance will increase relative to each initial population. As illustrated in Figure 1D, this will cause the risk probability distribution to widen, leading to increase in the risk of low-prevalence diseases, with no change at all for diseases with a prevalence of 50%.

The results of these simulations suggest that the true causal SNPs of polygenic diseases may easily differ by more than 30%, perhaps even by up to 100%, between geographically stratified populations, and clinical or epidemiological observations will be unlikely to register small and gradual increases in disease prevalence over successive generations because of the increase in the combined variance of a large number of risk alleles. A simulation of accelerated recombination with 1000 recombinations per parental generation genome resulted in the equilibrium level being reached within a few generations with a maximum prevalence increase of 45% when all SNPs differed from those in the original population and 6.3% when one-fifth of the SNPs differed. However, it would take many generations to reach this equilibrium, and on such a timescale, this process is likely to be indistinguishable in clinical practice from ongoing admixture with other populations and confounded by genetic drift, mutations, selection, stratification, environmental, and lifestyle changes.

### 2.2 Admixture of Populations with Differing PRSs

This scenario evaluated the admixture of two populations with similar polygenic EMOD architectures, where the higher-risk Population 2 was characterized by a common frequency of a small subset of alleles that had a very low frequency in Population 1, giving Population 2 an average relative risk (RR) of 10.0 (PRS difference in units of log(RR) = 2.30), as displayed in Figure 3. Accordingly, the initial disease prevalence was equal to 0.1% for Population 1 and 1% for Population 2. As expected from the conclusions of the preceding section, the relative PRS variance between the two initial populations before admixture was just 1.1%, even with the 10-fold difference in disease risk between the populations. This population liability is almost exactly reflected in Figure 1C, but not Figure 1D. The PRS effect size after admixture settled at the average between the two original populations, as is typical of the observational reports cited in the Introduction. The variance level of the combined population stabilized closer to the variance of the higher-risk Population 2, as would be expected from Equation (A1), with a negligible effect on the disease prevalence.

**Figure 3.**
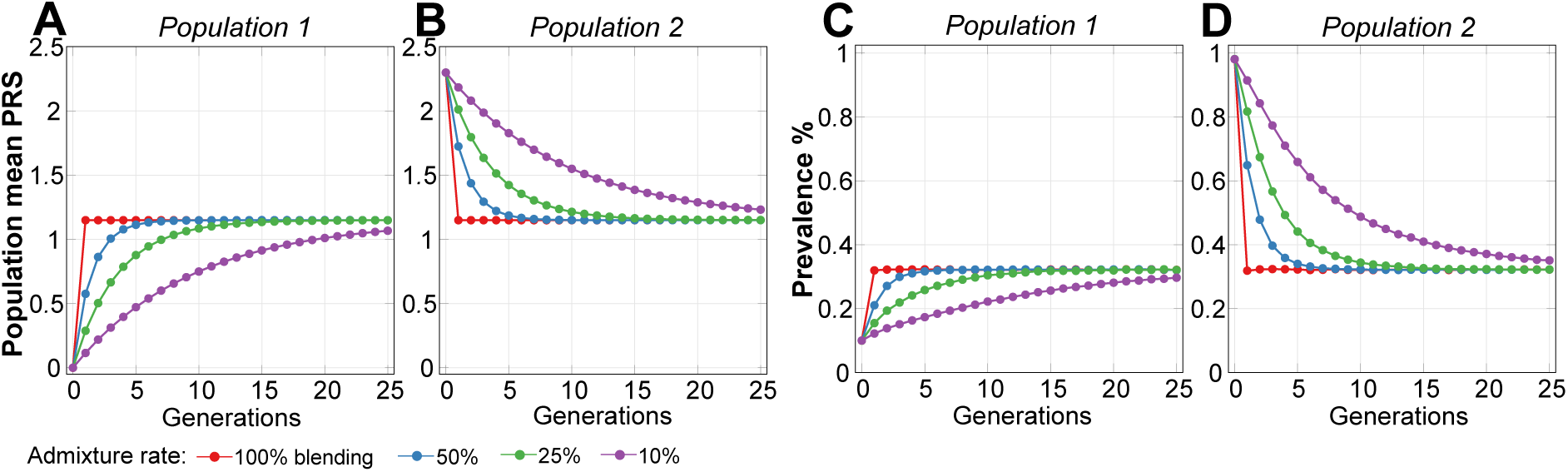
Admixture of two populations with a 10-fold relative risk difference. The plots (**A–B**) show the population mean polygenic risk score (PRS) equalizing between two populations depending on the admixture rates for a disease with 50% heritability: (**1**) shows Population 1 and (**2**) shows Population 2. Population 1, which was used as the reference, had a mean PRS of 0.00 and an initial prevalence of 0.1%. Population 2 is the higher-risk population and had an initial PRS of 2.30 and an initial prevalence of 1%. The plots (**C–D**) show the corresponding population prevalence change. Figure A2 shows a graphical display from the simulation and illustrates the admixture between these two populations.

Figure 3(A–B) show that the normalized PRS effect size difference between the populations accounted by the simulation almost exactly follows the proportion of population mixing under all admixture scenarios. This behavior matches the reported polygenic disease risk averaged in proportion to the population admixture noted in the publications referenced in the Introduction. While the admixture of two equally sized populations results in a precisely averaged PRS, the prevalence after mixing is close to the geometric mean of the initial prevalence values, resulting in a smaller-than-arithmetic average of the prevalence values of the initial populations. Thus, in this example, the prevalence is 0.32% rather than 0.55% (see Figure 3(C–D)). The PRSs will generally equalize following a simple mixing equation; this is true for both EMODs and LODs, as follows:

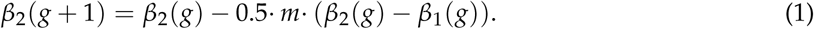

In the calculation for Population 2, *β*_2_(g) is the effect size (PRS) of Population 2 in generations *g* + 1 using values from the previous generation *g*. In this case, 0.5 is the ratio for equal population sizes, m is the admixture proportion, and *β*_1_ is the effect of Population 1; the equation for Population 1 mirrors Equation (1).

It is interesting to note that, even with a low 10% population admixture rate, the non-participating Population 2 prevalence decreases, relative to the baseline, to 91% in one generation, 84% in two generations, and 77% in three generations, and the improvement is even faster at higher admixture rates, with both populations heading toward an asymptotic admixed prevalence of 32%. Prominently, the equalization is reached in one generation in the 100% blending scenario (shown by the red lines in Figure 3).

### 2.3 Lowering Polygenic Disease Prevalence by Editing Effect SNPs

The gene therapy operations would change detrimental SNPs frequency in some fraction of a population. The population-wide Hardy-Weinberg equilibrium will be reached after one generation of random mating in an indefinitely large population with discrete generations, in the absence of mutation and selection, and the frequency of genotypes will remain constant across generations [68,69]. In case of high heterogeneity in effect alleles between populations, it may take a number of generations for the allele distribution to homogenize, accompanied with increase in disease prevalence, as was described in section 2.1. This effect is barely detectable for smaller risk allele differences, as modeled in the previous section 2.2.

Simulations confirmed that modifying or turning off a number of causal alleles in a higher-risk population can easily reduce the risk to that of a lower-risk population. Additionally, treating, for example, half of the individuals in a population with double the number of corrected SNPs (or any other proportion, as long as there are enough SNPs to correct) produces the same population risk load reduction, as the corrected SNPs would distribute within a few generations of random mating. Figure 4 demonstrates this by starting with a homogeneous population with identical risk in generation 0, subdividing individuals into two equally sized populations, and lowering the average RR of Population 1 by 10-fold (PRS = −2.3). The result is equivalent to those described for Population 1 and Population 2 in the previous section 2.2, as shown in Figure 3, and is followed by an identical admixture pattern. The variance of the combined population after admixture diminishes by 0.9%, reflecting the lower frequency of the risk alleles in the population.

**Figure 4.**
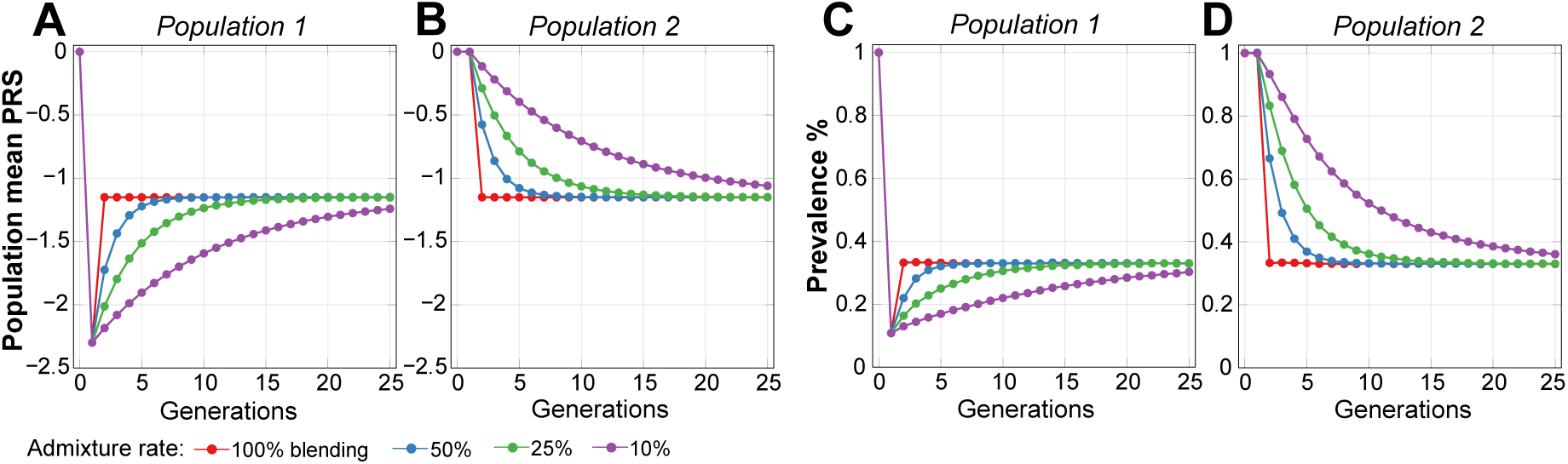
The application of gene therapy Population 1 lowers the population’s average risk by 10-fold, followed by admixture with Population 2. The homogeneous population is divided into Populations 1 and 2. In Population 1, an individual’s SNPs are uniformly edited to achieve a 10-fold improvement in relative risk (RR) (PRS = −2.30) in generation 1. The initial disease prevalence was set at 1%, and the disease heritability was 50%. Population 2 remains at this level in generation 1, while the prevalence of Population 1 decreases to 0.1%. After that, admixture patterns become mirror images of those in Figure 3. The plots in (**A–B**) show that the population mean polygenic risk score (PRS) equalizes between the two populations, depending on the admixture rate. The plots in (**C-D**) show a corresponding change in the mean population prevalence.

### 2.4 Estimates of Population Genomic Parameters for Diseases Known to have Large Risk Differences Between Ethnic Groups

Many diseases differ in terms of their risk and prevalence among subpopulations. In reviewed published cases, admixed populations were shown to have intermediate liability. Examples include differences in nicotine metabolism between Maori and European populations [39], differences in type 2 diabetes (T2D) risk between European American and African American populations [70], and differences in atrial fibrillation risk among a variety of populations [71], with prevalence usually differing by less than 2-fold between affected populations.

Three examples of diseases with contrasting risk between populations, primarily for middle-age-onset, are Dupuytren’s disease (DD), rheumatoid arthritis (RA), and lupus erythematosus (LE). DD heritability was determined by Larsen *et al*. [72] as 80%, with extremely varied prevalence, affecting at older ages 22-32% of men in populations originating from Northern European countries [57], and significantly lower prevalence in populations from other origins, with the lowest prevalence in Korea [73], Taiwan and China [74] at 100–1000 times lower prevalence than in Northern European populations. According to Molokhia and McKeigue [75], West Africans have a higher risk of LE than Europeans, and Native Americans have a higher RA risk than Europeans. Both diseases also show intermediate risks in admixed populations. LE heritability is estimated to be 44% [59,76], the prevalence was reported to be 0.35% for 60-year-old African American women and 0.1% for European American women [77]. RA heritability is estimated to be 60% [58], it has a prevalence of 3% in Canadian Native Americans and 0.3% in Europeans [78].

The above three examples were specifically chosen because their maximum incidence rates occur in early to late-middle ages. Therefore, prevalence of the diseases at moderately old ages approaches the disease lifetime risk. The admixture simulation results are presented in Table 2 and graphically illustrated in Figure A3.

**Table 2.**
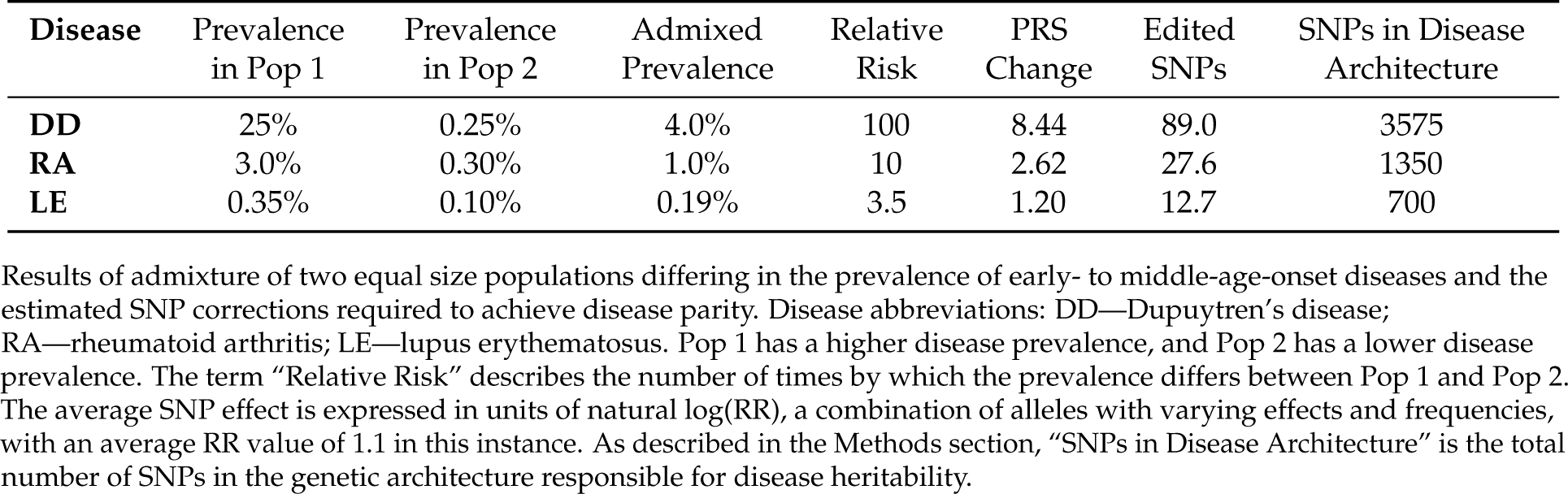
Admixture of diseases known to have large risk differences between ethnic groups.

| Disease | Prevalence<br>in Pop 1 | Prevalence<br>in Pop 2 | Admixed<br>Prevalence | Relative<br>Risk | PRS<br>Change | Edited<br>SNPs | SNPs in Disease<br>Architecture |
| --- | --- | --- | --- | --- | --- | --- | --- |
| DD | 25% | 0.25% | 4.0% | 100 | 8.44 | 89.0 | 3575 |
| RA | 3.0% | 0.30% | 1.0% | 10 | 2.62 | 27.6 | 1350 |
| LE | 0.35% | 0.10% | 0.19% | 3.5 | 1.20 | 12.7 | 700 |

Results of admixture of two equal size populations differing in the prevalence of early- to middle-age-onset diseases and the estimated SNP corrections required to achieve disease parity. Disease abbreviations: DD—Dupuytren's disease; RA—rheumatoid arthritis; LE—lupus erythematosus. Pop 1 has a higher disease prevalence, and Pop 2 has a lower disease prevalence. The term “Relative Risk” describes the number of times by which the prevalence differs between Pop 1 and Pop 2. The average SNP effect is expressed in units of natural log(RR), a combination of alleles with varying effects and frequencies, with an average RR value of 1.1 in this instance. As described in the Methods section, “SNPs in Disease Architecture” is the total number of SNPs in the genetic architecture responsible for disease heritability.

The last three columns in Table 2 show the differences in PRSs between populations (in units of log(RR)) and the average number of SNPs at the average genetic architecture effect size in need of correction to match the risk in high-risk populations with that in lower-risk populations if such a therapy were possible. It is shown that DD would require 89 SNPs to be corrected to reduce the high risk in North European ethnicities to match that in the Korean population. RA and LE would require significantly fewer edits. In each case, the number of edits constitutes only a small fraction of SNPs in each disease’s common low-effect genetic architecture.

The values of the admixed prevalence of RA and LE closely follow the geometric mean of the initial populations, as established in section 2.2. The simulation results noticeably deviate from the geometric mean in the case of DD, for which the geometric mean 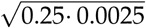 equals 2.5%, rather than the value of 4% found by the simulations. This indicates that 25% can hardly be considered a low prevalence from the perspective of relative risk, particularly when considering large risk differences between populations. Further simulation of scenarios with more common lower differences in disease relative risk between populations showed that prevalences after admixture closely followed the geometric mean of two initial populations; however, based on the assumption in Methods, the model is better confined to prevalences in single digits and below, typical to EMODs.

### 2.5 An Estimate of Preventive Gene Therapy for Early- to Middle-Age-Onset Polygenic Diseases

The review of the three diseases above—DD, RA, and LE—estimated the differences in the number of SNPs related to disease risks in naturally occurring populations and, accordingly, differences in the number of SNP corrections that would be required to achieve population parity for these EMODs.

Following the evaluation of population stratification by disease risk, admixture, and a simple correctional edit followed by population admixture in sections 2.2 and 2.3, it is time to consider a scenario that could allow for broader extrapolations. There can be countless potential scenarios of therapy levels, stratification, and admixture. It can be hypothesized that there may be an optimal level of population EMOD risk that can be achieved by lowering the average population PRS or, equivalently, by lowering the true causal risk allele frequencies.

A scenario was chosen in which, for the individuals participating in gene therapy (Population 1), the required number of risk SNPs was therapeutically edited to lower the population relative risk by 10-fold, or by a PRS of *β* = −2.3, in the first generation of ongoing therapy, on the premise that a 10-fold risk reduction in any disease would be a commendable improvement. Subsequently, smaller therapeutic interventions were applied in each generation to maintain Population 1 at this optimal level; the number of edits per generation is shown in Figure A4.

The evaluation of the admixture scenarios for Population 2, which does not directly participate in gene therapy (see in Figure 5), shows that in the 100% admixture (blending) scenario the disease prevalence in Population 2 to plummets to 0.32% (or 32% of the prevalence baseline value), while the population PRS reaches the exact halfway point between values in the original populations. However, unlike the admixture scenarios presented in sections 2.2 and 2.3, the improvement continues to asymptotically progress toward the treated Population 1 level of 10% of the baseline disease prevalence. The PRS progression using Equation (1) would just require fixing *β*_1_(*g*) = *Const*—the level of the chosen optimal treatment. From the perspective of the PRS admixture, this result is equivalent to the basic island-continent migration model; however, the disease prevalence connotations are noteworthy. Figure A5A also shows the renormalization of the relative PRS that can be applied to estimates with any chosen initial values of relative risk improvement, and in Figure A5B the normalized prevalence progression in case of the RR=10 treatment level. For comparison, the therapy alleviating population relative risk 4-fold depicted in Figure A5C showed that the relative prevalence reduction for the non-participating populations with ongoing admixture, as compared to the treated population, would be similar for varying degrees of treatment.

**Figure 5.**
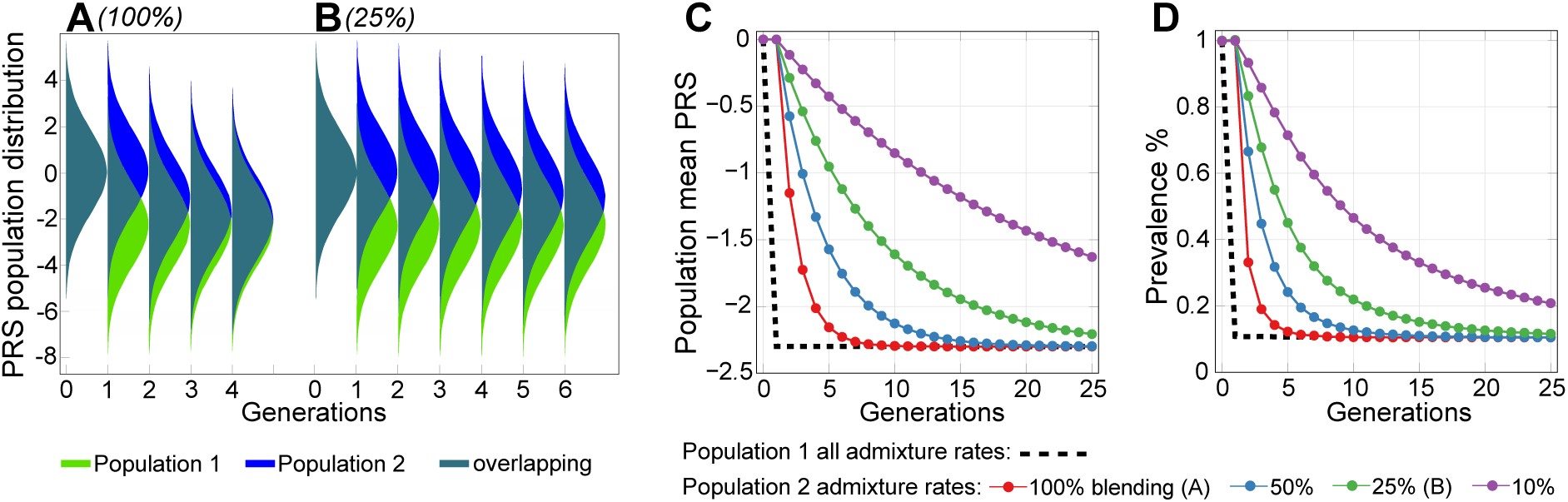
Preventive gene therapy with a 10-fold relative risk correction. A constant level of PRS = −2.3 is maintained for Population 1, undergoing admixture with Population 2. The 90° -rotated Gaussian-looking fills in plots (A–B) represent the population density for each generation at the corresponding PRS values in log(RR) units on the y-axis, and the colors represent the fraction of each population mix at each PRS value. (**A)** shows a 100% blending admixture, where the individuals from Population 1 mate exclusively with individuals from Population 2; (**B)** shows a 25% admixture, where individuals from each population have a 1/4 chance of of mating with individuals who are outside their own population; (**C**) shows that the population mean polygenic risk score (PRS) equalizing between the two populations, depending on the admixture rate; (**D**) shows the corresponding mean population prevalence change.

## 3. Discussion

With the accumulation of scientific knowledge of the genomic causes of common diseases and the advancement of gene-editing technologies, gene therapies to prevent polygenic diseases may soon become a reality. Over a decade of GWAS research has determined that polygenic EMODs and LODs share a genetic risk architecture: their causality is primarily attributable to common low-effect alleles [4,30] in multiplicative joint associations with environmental risk factors [31]. With the application of the multiplicative genetic risk model, the computer simulations developed in this research mapped the polygenic risk of the model genetic architecture of EMODs based on their prevalence and heritability. The results of these simulations correlated well with epidemiological observations (see Addendum A.1). Simulations of the admixture between modeled populations using this framework were performed to investigate a hypothetically possible range of heterogeneity of causal SNPs in geographically distinct populations. Subsequently, these simulations were applied to model scenarios of gene therapies to assess the relationship between population admixture and disease prevalence throughout generations. The simulations of admixture with differing causal SNPs between populations with identical disease prevalence demonstrated that, in principle, a large degree of heterogeneity in causal allele sets for EMODs between populations is possible. Whether all causal SNPs were identical or whether a large fraction of them differed between a pair of populations, the epidemiological and clinical statistics would be practically indistinguishable. Equally, it was shown that the outcomes of gene therapies would not be impeded under either situation. The commonality of causal gene variants for polygenic diseases between geographically distinct populations, as reported by GWASs [36,37,79] (with some models exploring a larger extent of allelic heterogeneity [80]), makes this extreme difference in causal allele sets unlikely, and the differences in disease prevalence and disease manifestation between populations appear to be primarily caused by differences in common allele frequencies. The finely balanced risk of genetic architecture in this model scenario would be far exceeded by the actual risk differences in geographically distinct populations, which often differ in disease prevalence [81]. The simulated population admixture for all polygenic diseases with differing risks among populations resulted in arithmetic averaging of the PRS, expressed as the sum of logarithms of the causal alleles’ true relative risk, and the prevalence of EMODs followed the geometric mean of the original populations.

The extreme differences in common EMOD risk, exemplified by DD, LE, and RA, demonstrate the range of polygenic distribution differences that may develop between populations because of geographic separation that occurs within an evolutionarily short time. Furthermore, these differences indicate the potential to alleviate these and other polygenic disease risks using gene therapy. The simulation results for typical EMODs show that the disease prevalence decreases in proportion to the degree by which the treatment lowers the population average relative risk.

It is hard to imagine that, even if such gene therapies were available, everyone would participate. In the hypothetical scenarios in which populations admix at a low rate of 10%—which would not be typical, particularly in the Americas [81]—the prevalence rates of the targeted diseases in the fraction of the population not directly receiving gene therapy would noticeably decrease in the second generation and even more so in subsequent generations. Longer term, this admixture would lead to a lower and more equal disease risk for all populations. A hypothetical example of such group stratification with regard to preventive gene therapy is preventive genetic treatment during in vitro fertilization (IVF), which could be legislatively limited only to situations in which the parents were found to possess high PRSs of a polygenic disease [14]. In the first generation, only the direct recipients would benefit, but normal admixture over the scale of generations would cause the whole population’s disease prevalence to diminish, as the simulations in this research demonstrate.

Again, hypothetically, even if gene therapy were to be discontinued after significantly reducing the risk of Mendelian diseases and EMODs over time, the low human germline mutation rate (estimated to be an average of 1.18 · 10^−8^ mutations per nucleotide per generation, which corresponds to 44–82 mutations per individual genome with an average of only one or two mutations affecting the exome [82]), means that many generations would pass before the disease rates would significantly increase again [83–85].

A complete picture of polygenic disease prevention must include LODs. The analysis method applied to EMODs would not be valid for polygenic LODs, because LODs typically manifest with extremely low incidences of diagnosis at younger ages, followed by a period of a nearly exponential annual increase in the disease incidence rate starting at relatively older and LOD-specific ages [50]. According to Chatterjee *et al*. [31], the conditional age-specific incidence rate of the disease can be modeled using Cox’s proportional hazards model [51] and multiplicative joint associations between low-penetrant SNPs and environmental risk factors [31]. An evaluation using this model [46] showed that a moderate level of therapy that lowered the hazard ratio by 4-fold (OR = 0.25) by converting detrimental SNPs to a neutral state would result in a delayed onset curve of LODs, with a delay of about 3 years for AD; close to 10–15 years for T2D, cerebral stroke, and coronary artery disease (CAD); and an even longer onset delay for breast, prostate, colorectal, and lung cancers.

A recent clinical and GWAS analysis by Mars *et al*. [52] determined that the difference in age at disease onset between the top and bottom 2.5% fraction of PRSs was 6–13 years for four LODs that overlapped with Oliynyk [46]. A lower onset difference value was found to be characteristic of T2D and CAD, while breast and prostate cancers showed the highest differences in terms of age of onset, thus clinically confirming the patterns predicted by simulations in [46]. The naturally occurring difference in the age of onset for the top and bottom fractions of the natural PRS variation [52], in principle, shows that applying gene therapy that would turn a sufficient number of true causal SNPs into neutral SNPs, thus turning the high risk population into the low risk population, would have the predicted outcome reflected in years of a delayed LOD onset.

The current research confirms that for polygenic diseases, including LODs, if gene therapy were to lower the frequency of true causal risk alleles and the corresponding population PRS, these proportions would propagate throughout subsequent generations [69]. In the case of admixture with populations not directly participating in gene therapy, the PRS would distribute proportionately to population mixing ratios, which for LODs will be reflected in disease onset delay [46] for all beneficiary generations. The incidence of EMODs does not strictly stop at a particular age; rather, a later but lower disease incidence occurs for all EMODs referenced herein. Therefore, preventive genetic treatment of these conditions may to a degree result in a delay of disease onsets.

In conclusion, the simulations in this research demonstrate that even if relatively large heterogeneity in the causal allele set for EMODs existed between populations, it will not be easily detectable by epidemiological studies in admixed populations. While the simulation results show that a large heterogeneity would be hypothetically possible, GWAS findings indicate the existence of a discernible commonality of causal SNPs for polygenic diseases between geographically distinct populations, and the extent of the risk differences between populations due to unique causal SNPs is likely not extreme. Even if it were large, this potential difference would not impede the outcomes of preventive gene therapies if they were applied to turn population-specific true causal SNPs to a naturally existing neutral state of nucleotides, and this would hold after populations admix.

Preventive gene therapy that is designed to turn true causal SNPs into a naturally existing neutral state of nucleotides would result in a decrease in EMOD prevalence proportionate to the decrease in the population relative risk attributed to the edited SNPs. The outcome will manifest differently for LODs, where the therapies would result in a delay in the disease onset and decrease in lifetime risk, however the lifetime risk would increase with prolonged life expectancy, a likely consequence of such therapies. EMODs exhibit some degree of incidence later in life, and hypothetically, some of the outcomes may share characteristics with LODs.

In summary, the results of this study show that if gene therapies of preventive heritable diseases were to be applied on a large scale, even with a fraction of the population participating, the decreasing frequency of risk alleles in the population would lower disease risks or delay the ages of disease onset. With ongoing population admixture, all groups would benefit throughout successive generations.

## 4. Methods

This study assessed population genetics dynamics for a hypothetical future in which gene therapy can be applied to prevent polygenic diseases. In earlier research, the risk allele distribution for polygenic LODs that accompanies aging was evaluated [50], and the potential of future preventive gene therapy to delay onset ages and lower the lifetime risk of developing such LODs was successfully quantified [46], as demonstrated in Figure 1B, by leveraging age-specific incidence rates under multiplicative [31] Cox’s proportional hazards model [51]. The findings of this earlier publication complement the results of the current research and are noted in the Discussion.

The main goal of this study was to quantify the impact of gene therapy from a population genetics perspective while accounting for population stratification and admixture. The gene therapy corrections that change detrimental SNPs frequency within a subset of a population, will reach population-wide Hardy-Weinberg equilibrium after one generation of random mating in an indefinitely large population with discrete generations, in the absence of mutation and selection, and the frequency of genotypes will remain constant throughout generations [68,69]. This equally applies to polygenic phenotypes [86], and the extended diploid Wright–Fisher model simulation reproduced this expected behavior, thus validating that the model’s granularity on a generational scale was appropriate for the intended target of this research. Although the mean population PRS found in this study precisely follows the Hardy–Weinberg principle, the behavior of disease risk variance in the polygenic admixture is more gradual as a result of linkage disequilibrium and recombination [67,87,88].

The following sections review the simulation’s conceptual foundations and conclude by describing the simulation steps.

### 4.1 Considerations for Liability Threshold Models

Of the polygenic diseases analyzed in this research, those with the highest incidence in early- and middle-age are less suitable for the age-specific rates approach used earlier for LODs [46], because subjects with an earlier age at onset do not necessarily show an increased polygenic risk burden, as exemplified by the incidence of schizophrenia [64]. The prevalence of these diseases is sometimes modeled using the liability threshold model, originally proposed by Falconer [47 48]. Under this model, illustrated in Figure 1(C–D), the disease prevalence is a function of disease liability, which is represented by polygenic risk. In the liability threshold model, an individual can be characterized by a genetic liability to a disease. A combination of genetic and environmental effects results in a probabilistic disease distribution among individuals. In the original Falconer [48] interpretation, all individuals whose PRS exceeds the threshold contribute to the disease prevalence; graphically, these individuals fall to the right of the threshold. Subsequent research has shown that the multiplicative risk model is most suitable for explaining experimental data. This model is exemplified by three approaches: the Risch risk model, the odds risk model, and the probit risk model [49,89,90]. The solutions based on these models are typically obtained through simulations or numerical methods, with the exception of the simplest scenarios that allow for analytic solutions, providing estimates of disease prevalence according to the polygenic risk distribution. These models lack the ability to sample individuals in the multi-generation population simulations required in this study, and they are also based on specific allele distributions that will not be maintained during ongoing admixture and gene therapy. Hence, this study developed the simulation approach described in section 4.4, applying probabilistic sampling of individuals by PRS validated in [46].

### 4.2 Conceptual Summary

The simulated diseases were assumed to have an early- to middle-age onset, with a negligible disease incidence at older ages. The term “prevalence” is customarily used in liability threshold models. However, often, whether the term pertains to a whole population or a population of a certain age range is not well defined. Herein, the term is used in a narrower scope; in this study, “prevalence” means the cumulative incidence of a disease at an age later than the typical onset age range, with negligible incidence later on. Thus, the definition of prevalence in this context is more similar to the lifetime risk concept.

The heritability of EMODs usually ranges from 30% to 80%, as documented by Wang *et al*. [65] and Polubriaginof *et al*. [66]. A heritability level of 50% was chosen for most simulations and analyses to represent a typical EMOD, and the common low-effect-size genetic architecture SNP set was assembled accordingly, as noted in section 4.3. The analysis of specific EMODs used their heritabilities.

Large population sizes were used to make genetic drift effects imperceptible at the short generational scale used in the simulations. Similarly, although the simulation design allowed for the introduction of mutations, given the short generational scale under consideration, mutations could not achieve common population frequency, [83–85] and were not introduced.

This study was not concerned with evaluating potential obstacles due to pleiotropy, which, in the context of gene therapy, is defined as the possible negative effects on other phenotypic features resulting from an attempt to prevent an EMOD by modifying a subset of SNPs [91,92]. Under the common low-effect genetic architecture used in the simulations, from an average of 514 such SNPs in the average modeled individual, gene therapies would only need to correct an average of 15 SNPs to achieve a 4-fold decrease in the relative risk (PRS = −1.386) and 24 SNPs to achieve a 10-fold RR decrease (PRS = −2.30), as shown in Figure A1A. Arguably, with personalized prophylactic treatment, it would be possible to select a small fraction of variants from a large set of available choices, as exemplified in Table 2, that do not possess antagonistic pleiotropy, or perhaps even select SNPs that are agonistically pleiotropic with regard to some of the other EMODs and LODs. After all, because of a balance between selection, mutation, and genetic drift on evolutionary scales [84], a proportion of low-effect detrimental SNPs have achieved common population frequency, simply because they were not detrimental enough to have been selected out, rather than having been selected for because they provide a physiological or survival benefit. Thus, these SNPs would constitute an uncontroversial therapeutic target.

In the simulations, the F-statistic (Fst) for disease architecture alleles was calculated using Hudson’s method, as recommended by Bhatia *et al*. [93], and the alternative allele frequency difference (AFD) statistics were also calculated [94]. The statistics obtained were unsurprising for the simulated populational processes, and including their interpretation in the reported results would be extraneous. Nevertheless, for those interested, these results are available in Supplementary Data. While admixture naturally involves multiple world populations, simulating the admixture of two populations was adequate for the intended analysis and extrapolations.

The analysis in this study is contingent on future genetic and computational techniques being capable of determining and safely modifying a relatively small subset of disease genetic architecture SNPs from a detrimental state to a neutral one. This is easy to accomplish in a population simulation, in which the effect sizes and states of detrimental SNPs are known for each individual. These model genetic architecture SNPs are treated as variants that are truly causal for disease liability and heritability. A brief summary of current gene-editing technologies is included in Addendum A.2.

### 4.3 Allele Genetic Architecture

The common low-effect-allele architecture was implemented in a similar manner to that used in the author’s earlier research [50], which followed the approach used by [17]. The summary, including specifics of the implementation in this study, is available in Addendum A.3. In contrast to GWAS tag SNPs, the model genetic architecture SNPs are truly causal for disease liability and heritability variants, and they are assumed to be accurately identified for the purposes of personalized gene therapy. Estimates using the liability threshold model customarily use RR values to model known causal SNPs [49,95]. This research followed suit: SNP effects were treated in terms of relative risk, and PRSs were expressed in terms of the sum of the logarithm of RR. This method is also justified by the fact that the majority of EMODs have a prevalence of less than 2%, as exemplified by RA [58], LE [59], schizophrenia and bipolar disorder [60,96,97], and Crohn’s disease [61,62], with only a small number of diseases such as asthma [53] approaching a prevalence of 10% [65]. Dupuytren’s disease, which has a prevalence of more than 30% in some Northern European ethnicities, although it is lower in most of the world by 1–3 levels of magnitude, is an interesting example that was examined in this research. The alleles were randomly distributed throughout the model genome; these results are consistent with GWAS findings for asthma [53,98], schizophrenia [99], and other diseases [4].

### 4.4 Disease Prevalence Analysis

In order to track the changes in disease prevalence associated with population admixture and gene therapy, it was necessary to map PRSs to the probabilities of succumbing to a polygenic disease on the basis of the genetic architecture and disease prevalence. Individual RRs *R*_*i*_ were calculated as a product of the RRs of all SNPs in the disease genetic architecture, as follows:

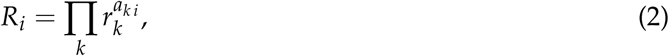

where *r*_*k*_ is the *k*th SNP’s true RR, and *a*_*k i*_ (equal to 0, 1, or 2) is the number of the *k*th allele in a pair of individual chromosomes *i*. The PRS *β*_*i*_ = *log*(*R*_*i*_) is defined in Addendum A.3. Multiplicativity by RR is equivalent to additivity by PRS.

The simulations sampled individuals from the allocated population without replacement, proportionate to individual RR *R*_*i*_, until a sample size of *n* individuals—those diagnosed with the disease—reached the number that satisfied the disease prevalence *K*:

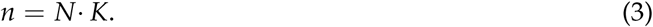

The goal was to map an individual’s PRS to the probability of them becoming ill on the basis of disease prevalence and PRS distribution, dictated by heritability and allele genetic architecture, as follows:

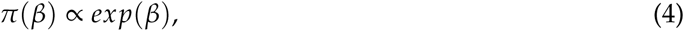

In practice, the simulation loop sorted the sampled diagnosed individuals into narrow PRS intervals, from *β* to *β* + Δ*β*, and determined the probabilities *π* of each PRS band, as follows:

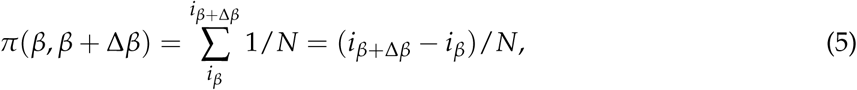

where *i*_*β*_ to *i*_*β*+Δ*β*_ are numbers of individuals sorted by PRS in a PRS band, and *N* is the population size.

Thus, under the multiplicative risk model, an individual’s probability of being diagnosed with the disease under consideration can be mapped to the individual PRS, and this mapping can be used in subsequent generations after gene therapy and population admixture. The advantage of this approach is that once the mapping is determined, it can be saved and reused in subsequent simulation runs as long as the chosen initial genetic architecture and prevalence are identical. This initial mapping was made very accurate by building large sets of individual PRSs per run of determination simulation (a set of eight billion was typically used) and averaging the mapping over multiple runs. The resulting mapping distribution is shown in Figure 6.

**Figure 6.**
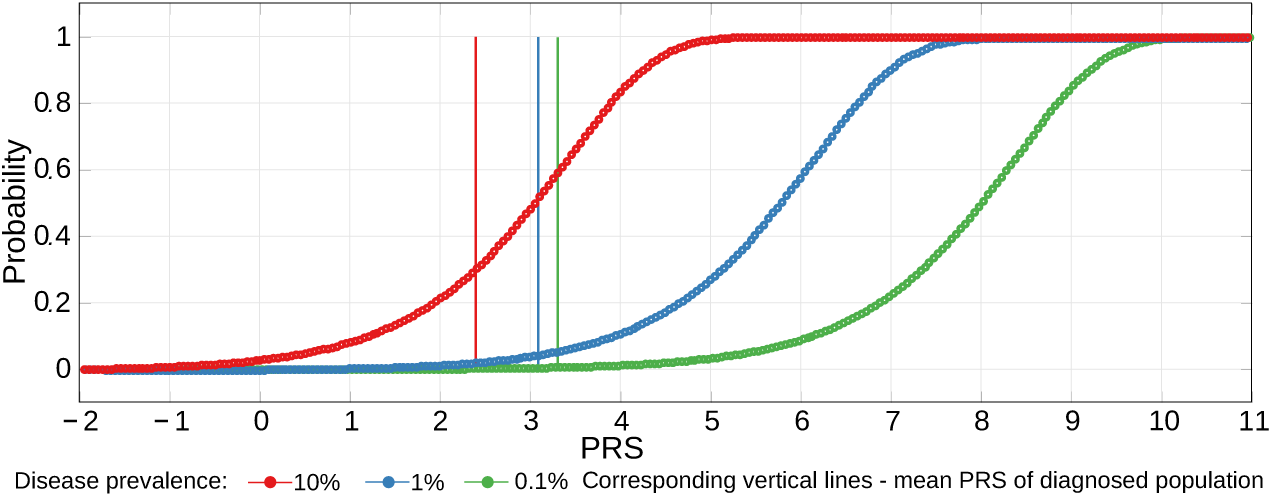
Disease probability distribution mapped to individual PRS. In simulations for a population with a mean PRS normalized to zero and a heritability of 50%, the PRS probability of disease curves reproduces the liability threshold model’s logistic distribution of probabilities [100]. This PRS probability distribution allows for the precise reproduction of the original disease prevalence and is used to determine changes in prevalence that result from simulated population admixture and gene therapy. The mean PRS of a diagnosed population and the probability curve move toward lower values, as illustrated in [49]. These mean diagnosed PRSs are not the liability threshold values because of the population density distribution being Gaussian with a zero mean and the probability curves represented in this figure.

The application of this mapping, using identical PRS bands, to the initial population reproduced the original prevalence with high precision and obtained a deviation of less than 2% in a two-sigma (95%) confidence interval for the PRS and prevalence results. Thus, error bars in the graphs would be extraneous. An exception is the population admixture figures in which a small relative change in values necessitated the inclusion of the two-sigma error bars (for example, in Figure 2C).

### 4.5 Simulating Gene Therapy Under Population Stratification and Admixture Scenarios

The following simulation steps were performed.

#### (1) Simulation initialization

The simulation initialization steps were performed, including the allocation of population objects and the assignment of individual PRSs on the basis of the modeled genetic architecture allele frequencies chosen for each population. Individuals were subdivided into two populations, Populations 1 and 2, with equal relative sizes and male/female proportions (configurable in the simulation setup). The initial disease prevalence and genetic architecture effect size in Population 1 were always used as references for Population 2 and the combined population. When gene therapy was performed, it was always applied to Population 1. For the validation of extreme population stratification and admixture scenarios, four sets of genetic architectures were constructed and specified in the simulation configuration. The population differences were set to 100% (all causal SNPs differ between population genetic architectures), 66%, 33%, and 20% (i.e., one-fifth of the causal SNPs differ). The difference was estimated by the fraction of the PRS difference that was attributed to differing SNP architectures between the two populations.

#### (2) Reproduction

The simulation proceeded through successive generations via reproduction with the configured level of population admixture. The admixture was configurable in a range from 100% to 0%. The rate of 100% meant that exclusively members of the opposite populations reproduce with each other (also referred as “blending”, where either population contributes exactly half of the diploid genome to each offspring in a generation). Above 50% the reproduction is preferentially between opposite populations. The 50% probability means that there is an equal probability that reproduction occurs within the same population and between opposite populations. Lower than 50% values, for example, an admixture level of 10% means that the probability of individuals reproducing within their own population is 90%, and the chance of admixture with the other population is 10%. The offspring of the opposite populations had an equal chance to belong to either population, and the offspring from reproduction within the same population remained in their parents population.

#### (3) Recombination

Because the parental pairs were chosen in the preceding step, each parent’s genome proceeded through recombination. The reported results used an average of 36 Poisson-distributed recombinations per parent in a single linear genome (configurable), and accelerated recombination of 1000 average Poisson-distributed crossovers was used to validate population admixture with a high level of difference in disease genetic architectures between populations.

#### (4) Gene Therapy

The gene therapy step consisted of sampling risk alleles for each individual chosen as a subject for gene therapy. The requisite number of risk alleles were turned off in order to achieve the chosen PRS improvement. As expected, the population average PRS reached equilibrium during the generation of random mating. The same PRS improvement was achieved by applying the same level of cumulative therapy to the highest-risk individuals or by averaging it over the population or any other population subset. Of the available simulation options, two were found to be the most illuminating: (a) therapy in a single generation of Population 1, followed by a varying degree of admixture with Population 2, and (b) the continuous maintenance of a chosen optimal population health improvement (PRS level) in Population 1, accompanied by varying levels of admixture with Population 2. Gene therapy included the ability to define the set of SNPs to be edited. This was carried out by specifying the desired SNPs in a configuration file, which was valuable for validating the results shown in section 2.2.

#### (5) Analysis

The individual risk alleles in each individual were accounted for at a number of stages in the simulation process and aggregated into the population PRS distribution, prevalence analysis, and Fst and AFD statistics, which were saved in comma-separated values format for further analysis and reporting.

#### (6) Repeat

Steps (2)–(5) were repeated until the defined generation limit was reached. The simulation flow configuration included the option of re-running the same simulations multiple times. This allowed the results of multiple simulation runs to be averaged and the resulting multi-run variance and standard deviation for key statistics to be determined.

The simulation configuration screen, which references the described and additional options, can be seen in Figure A6.

## Acknowledgments

The author thanks Alexei J. Drummond and Peter R. Wills at the University of Auckland.

## Abbreviations

The following abbreviations are used in this manuscript:

AD: Alzheimer’s disease
AFD: allele frequency difference statistic [94]
CAD: coronary artery disease
DD: Dupuytren’s disease
EMOD: **E**arly- to **M**iddle-age-**O**nset polygenic **D**isease
Fst: F-statistics, originally conceived as te fixation index by Wright, implemented here using Hudson’s method [93]
GRS: genetic risk score; used synonymously with polygenic risk score, abbreviated below
GWAS: genome-wide association study
LE: lupus erythematosus
LOD: late-onset disease; herein, analyzed LODs are exclusively polygenic
MAF: minor allele frequency; customarily implies the effect allele frequency
OR: odds ratio
PRS: polygenic risk score; in this study, a normalized sum of logarithms of additional relative risk conferred by causal alleles
RA: rheumatoid arthritis
RR: relative risk or risk ratio
SNP: single nucleotide polymorphism; in the context of this study, SNP is used synonymously with the term ‘allele’
T2D: type 2 diabetes
WGS: whole genome sequencing ff

## Appendix A. Supplementary Chapters and Figures

### Appendix A.1. Population Stratification and Admixture from the Perspective of Polygenic Disease Risk

Geographic and local population genetic stratification and variation complicate the ability to diagnose and treat a number of medical conditions [32]. It is well known that people from different geographic origins may have different rates of specific diseases, physiological responses to medications, and as a result, different medical treatment outcomes. For example, in the US, the prevalence of type 2 diabetes is 12.8% in African Americans, 8.4% in Mexican Americans, and 6.6% in non-Hispanic whites [38]. Belbin *et al*. [40] investigated the difference in allele frequencies among individuals in Latin American populations and found that although they were ostensibly derived from the same population, the top and bottom quartiles of the dominant ancestral component in admixed populations had larger changes in allele frequencies, with 20.4% of sites exhibiting a difference in frequency of >10% in individuals in the upper and lower quartiles with European ancestry in Puerto Rico. For individuals with Native American ancestry in a Mexican population, 36.0% of sites differed by >10%. This characteristic is shared by all groups that have undergone recent admixture and has been magnified by the multi-continental ancestry and local differentiation that underlie the genetic history of Latino populations [40]. A study that reviewed the US Veteran Affairs database [71] found age-adjusted prevalence values of atrial fibrillation (AF) of 5.7% in European Americans, 3.4% in African Americans, 3.0% in Hispanics, 5.4% in Native Americans/Alaskans, 3.6% in Asians, and 5.2% in Pacific Islanders. The differences in prevalence were accompanied by differences in AF symptoms, management, response to anticoagulants, and outcomes for these populations [101]. Another example is coronary artery disease genetics, which also varies in prevalence among populations [102]. In addition, breast cancer incidence is higher for Puerto Ricans and Cuban Latinas than for those from Mexico [103], and there are statistical differences by national origin in the rates of prostate, colorectal, lung, and liver cancers [103].

The predictive ability of GWAS and GWAS PRSs also varies broadly if the score is being applied to a population other than the one for which the score was initially determined [34]. For example, Holley *et al*. [104] observed significant differences in the distribution of SNPs associated with disease risk in New Zealand Maori patients with myocardial infarction compared with those of European origin. The authors concluded that although the genetic risk score (GRS) is overall higher for Maori when applying existing GRS tools, careful evaluation is needed before internationally developed GRS tools can be applied. Africa’s haplotype diversity, which is the highest on Earth, has important implications for the design of large-scale medical genomics studies across the continent [105]. Investigations by local research institutions, given their rich local clinical data and case-control base, could help to bridge the existing knowledge gap and provide valuable nuanced genomic information for these communities and their descendants, including those who have emigrated to other regions of the world.

At the same time, there are indications of commonality of gene variants that are causal for polygenic diseases among geographically distinct populations. A study by Seyerle *et al*. [36], which was performed for five geographically distinct populations, found that of 21 SNPs implicated as genetic determinants in QT-interval prolongation, seven showed a consistent direction of effect in all populations, and nine showed a consistent effect for four populations and typically small opposite effects for the remaining population. The effect allele frequency (EAF) varied among these populations. A GWAS on 28 diseases in Europeans and East Asians was conducted by Marigorta and Navarro [37], who reported high trans-ethnic replicability, implying common causal variants. Admixed populations usually show an intermediate level of liability or effect. For example, in individuals of Maori descent, nicotine metabolism is 35% lower than that in Europeans, with the metabolism of admixed individuals fitting between those of the two populations [39].

A simulation study by Zanetti and Weale [106] found that a combination of Euro-centric SNP selection and between-population differences in linkage disequilibrium and EAF was sufficient to explain the rate of previously reported trans-ethnic differences, without the need to assume between-population differences in the true causal SNP effect size. These findings suggest that the cross-population consistency found in this study is larger than that usually reported. Martin *et al*. [42] stated that, contrary to the belief that the polygenic scores of diverse populations are doomed to produce low PRS predictive power, diverse cohorts, rather than homogeneous cohorts, should be used. The authors further claimed that the effect size estimates from diverse cohorts are typically more precise than those from single-ancestry cohorts, and the resolution of causal variant fine-mapping can be considerably improved.

### Appendix A.2. A Concise Summary of Gene-Editing Techniques

This study analyzed the outcomes of simulated population genetics in response to hypothetical future gene-editing therapy for the prophylaxis of polygenic heritable diseases. Many ethical and regulatory considerations will need to be settled before (or if) such therapies become practicable [15,107]. Deeper scientific knowledge and more advanced techniques are being developed, particularly for personalized determination (with either computational methods or thoroughly verified genomic databases) of the deleterious effects of common [108–110] and rare allele variations and exome mutations [43,111–116]. It may be many years (likely decades) until precise knowledge is of sufficient depth for personalized medicine diagnostics to be conducted.

Gene-editing techniques need to perfect the ability to precisely modify genomics sequences with minimal off-target defects and to develop robust quality control measures for the edited results. The most promising current technology is CRISPR-Cas9 [117], a rapidly developing technology, which replaced older technologies such as zinc-finger nuclease (ZFN) [118] and transcription activator-like effector nuclease (TALEN) [119]. In 2019 alone, reports were published on the improved specificity of the CRISPR operation [120], the modification of thousands of nucleotides while reducing DNS nicking [121], and the use of CRISPR-associated transposons to insert custom genes into DNA without cutting it [122], among many other developments. Synthetic genomics, which is mostly in the proof-of-concept stage [123,124], could be another promising future technology. Continuous improvement is required for the technology to reach sufficient specificity, access all areas of the genome, and achieve a sufficiently low number of off-target edits and defects. Only following this will it be well suited for routine gene-editing therapeutic use.

### Appendix A.3. Implementation of Common Low-Effect Genetic Architecture

The genetic disease architectures used in this research were based on [17]—a simulation study that determined the number of alleles needed to achieve a statistical distribution variance that corresponds to the heritability of a particular polygenic disease or phenotypic feature. The allele architecture scenarios were implemented in the simulations in an identical manner to that applied in earlier research, in which five genetic architectures were validated and common low-effect-size genetic architecture was determined to indeed best fit the observed experimental and clinical data (see [50] for a comprehensive description). The common low-effect-size genetic architecture was used throughout this study. A concise summary of its major concepts, reformulated in terms of allele relative risks and the implementation steps that differed in this research follows.

The resulting variance of the allele distribution was determined to be

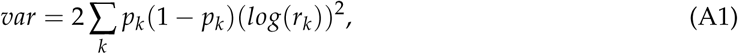

where *p*_*k*_ is the frequency of the *k*th genotype, and *r*_*k*_ is the relative risk of any additional liability presented by the *k*th allele for a particular individual. The contribution of genetic variance to the risk can be expressed as the disease heritability:

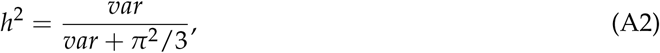

where *π*^2^/3 is the variance of the standard logistic distribution [125].

Following [17], the variants were assigned to individuals with frequencies proportionate to the minor allele frequency (MAF) *p*_*k*_ for SNP *k*, producing, in accordance with the Hardy–Weinberg principle, three genotypes (AA, AB, or BB) for each SNP with frequencies of 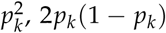, and (1 − *p*_*k*_)^2^. The diploid Wright–Fisher model simulation with recombination requires the tracking of SNPs on two chromosomes, and the individual PRSs *βind* for *k* SNPs were calculated as follows:

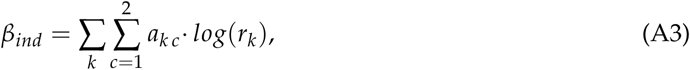

where *a*_*k c*_ (0 or 1) is the state of the *k*th SNP on chromosome *c* (1 or 2), and *r*_*k*_ is the relative risk of the additional liability presented by the *k*th allele for a particular individual. The population mean PRS value *β*_*mean*_ was calculated from the genetic architecture distribution using the following equation:

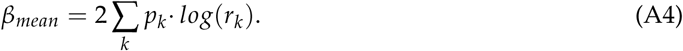

Two populations were always used in the simulations, and the *β*_*mean*_ value of the first (reference) population was applied to both populations, making it easy to compare the populations’ PRSs, as well as the distribution of higher- and lower-risk individuals within and between the populations.

For the common allele low-effect-size genetic architecture model, which, based on [17], was expected to be the most suitable for explaining the heritability of the analyzed LODs, the risk alleles were discretized into five equally spaced values within the defined range, with an equal proportion of each allele and an equal odds ratio in each. In this case, the MAFs were distributed in equal proportions of 0.073, 0.180, 0.286, 0.393, and 0.500, while the relative risk (RR) values were 1.15, 1.125, 1.100, 1.075, and 1.05. Thus, 25 combinations were possible. These entire blocks were repeated until the target heritability level was achieved, which, in this case, was 36 times for *h*^2^ = 50%. Figure A1 demonstrates the populations’ SNP and PRS distribution for the 50% heritability scenario and the 80% scenario used in the simulations of Dupuytren’s disease. The genetic architecture scenarios were defined in comma-separated values (CSV) files in the executable folder. In this study, the files were given names such as ‘A0.txt’ and ‘A11S.txt’.

**Figure A1.**
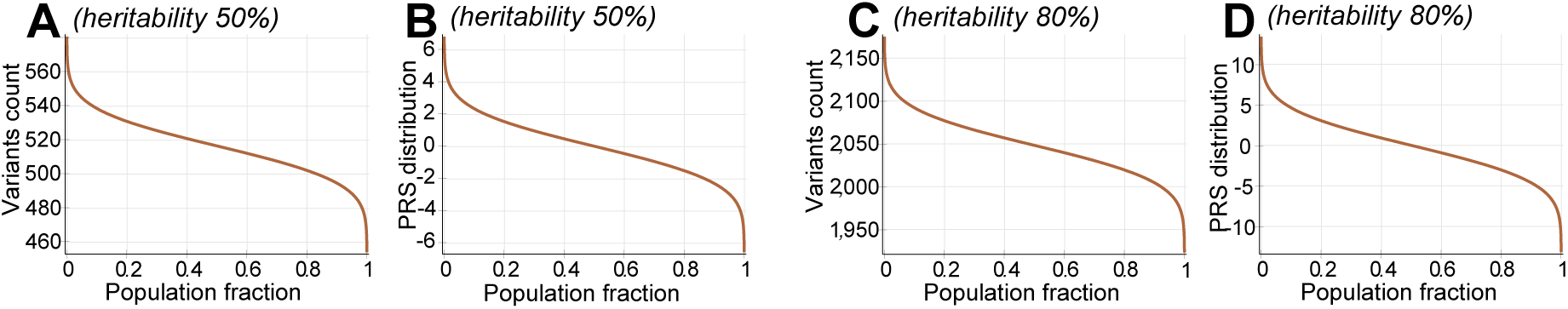
Population distribution of detrimental variant counts and polygenic risk scores (PRSs) for common low-effect-size genetic architecture. (**A**) Variant count for a simulated disease with 50% heritability; (**B**) the PRS for a simulated disease with 50% heritability; (**C**) variant count for a disease with 80% heritability, exemplified by Dupuytren’s disease (DD); (**B**) the PRS for DD, with 80% heritability.

The file *‘ConditionFiles.txt’* specifies which genetic architecture files were loaded for a given simulation run. It contains two columns: the first column defines the number of times to repeat the loading of a genetic architecture file to achieve the desired heritability, and the second column specifies the name of the genetic architecture file. The simulations always operated on two populations. Therefore, two files were always specified in two lines; the ‘#’ symbol was used as the first character in a line to comment out that particular line. Genetic architectures can be specified by the same architecture file when the initial population is homogeneous, or each file can represent different allele frequencies, but the effect sizes must match.

Only the following columns from the genetic architecture files were applicable to these simulations (the remaining columns were set to 0 and ignored): *SNP* denotes the RSxxx-style SNP identifier; *EAF* is the effect allele frequency; and *OR* is the odds ratio, which is actually the allele relative risk in this case. Additional SNP lines were used to facilitate specific analyses. This was accomplished by duplicating the entries and setting alternate entries to either the EAF required in the genetic architecture or a low frequency to simulate a different allele or an allele that was not represented in the populations.

### Appendix A.4. Supplementary Figures

**Figure A2.**
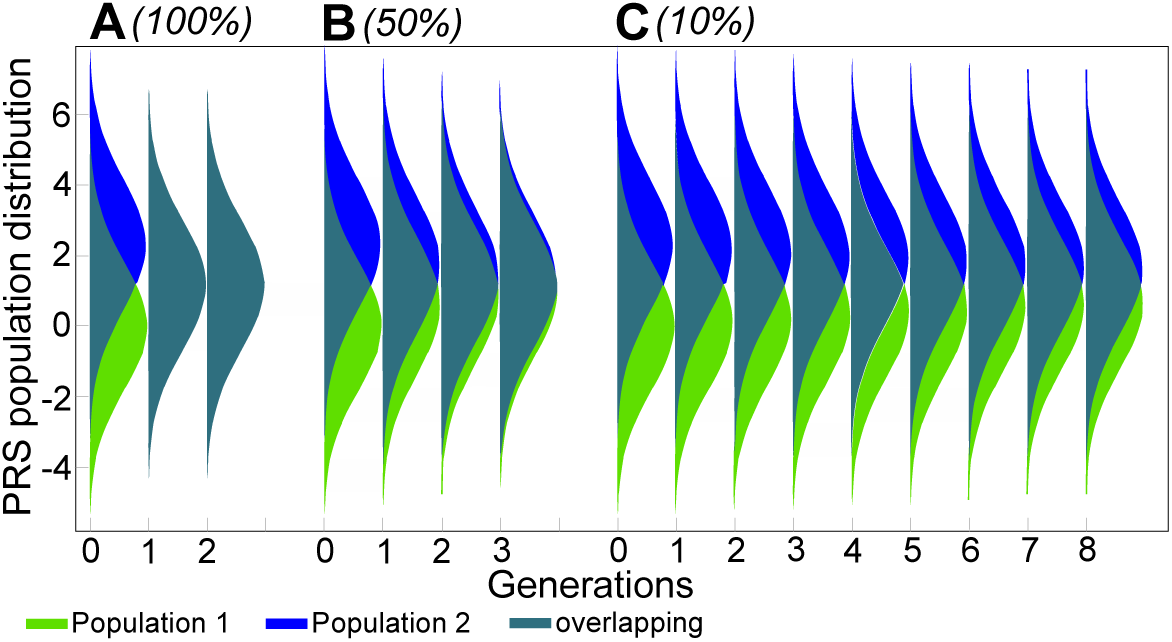
Graphical display from the simulation screenshot showing the admixture of two populations with a 10-fold difference in relative risk. The 90°-rotated Gaussian-looking fills represent the population density for each generation at the corresponding PRS values in log(RR) units on the y-axis, and the colors represent the fraction of each population mix at each PRS value. (**A)** shows the 100% blending admixture, in which the individuals from Population 1 mate exclusively with individuals from Population 2; (**B)** shows the 50% admixture, in which individuals from each population have an equal chance of mating within their population and with the other population; (**C)** shows the case in which individuals in Population 1 have a 10% chance of mating with individuals from Population 2 and vice versa.

**Figure A3.**
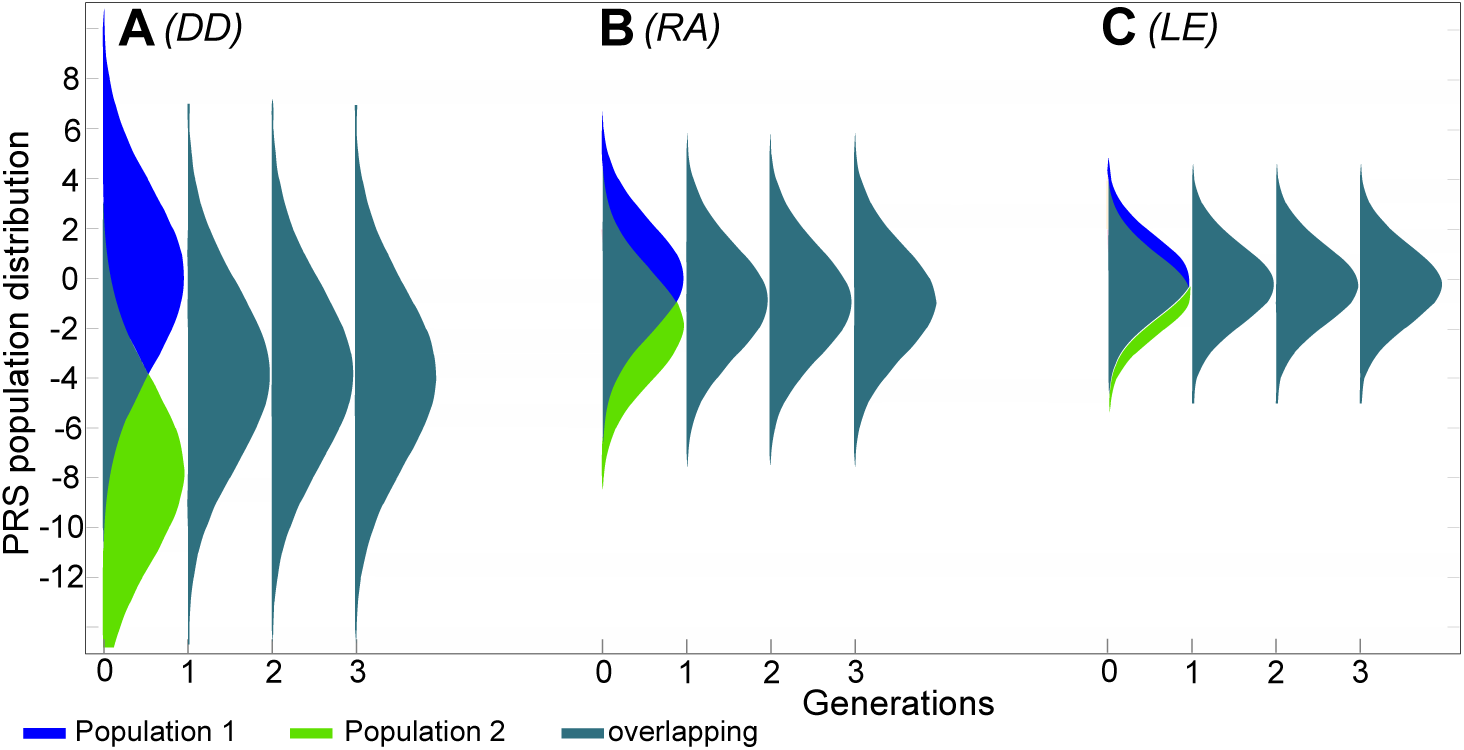
Graphical displays from the simulation screenshots showing the admixture of diseases that are known to have large differences in risk between ethnic groups. Population 1: higher-risk population (reference); Population 2: lower-risk population. The plots illustrate a scenario with 100% population blending between Populations 1 and 2, which are equal in size. The 90° -rotated Gaussian-looking fills represent the population density for each generation at the corresponding PRS values in log(RR) units on the y-axis, and the colors represent the fraction of each population mix at each PRS value. (**A)** Dupuytren’s disease (DD), heritability 80%, Population 1 prevalence of 25%, Population 2 prevalence of 0.25% (100 times the relative risk). (**B)** Rheumatoid arthritis (RA), heritability 60%, Population 1 prevalence of 3%, Population 2 prevalence of 0.3% (10 times the relative risk). (**C)** Lupus erythematosus (LE), heritability 44%, Population 1 prevalence of 0.35%, Population 2 prevalence of 0.10% (3.5 times the relative risk).

**Figure A4.**
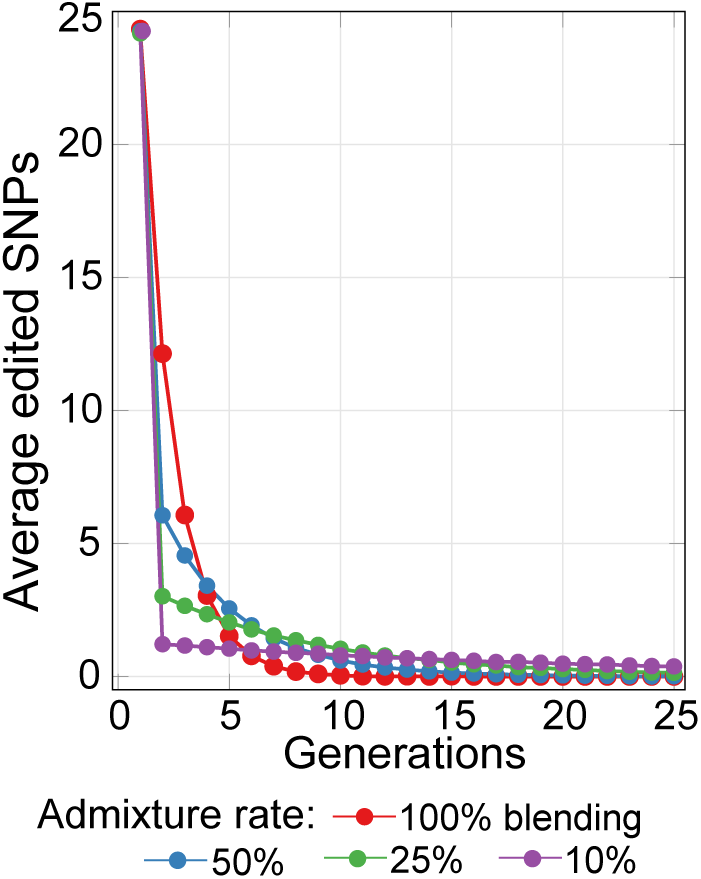
The average number of edited SNPs per individual in the scenario in which gene therapy maintains a constant optimal level of disease risk in the population participating in gene therapy, with differing degrees of admixture with a non-participating population. The first population depth of the edit was identical for all admixture scenarios. The highest admixture rate among populations led to a continually higher number of edits, and the asymptotic balance—the point at which maintenance edits were no longer needed—was reached more quickly. Comparatively, lower levels of admixture needed a lower initial number of additional edits per generation to maintain a constant risk level for the population participating in therapy. However, the number of generations required to reach equilibrium was much larger.

**Figure A5.**
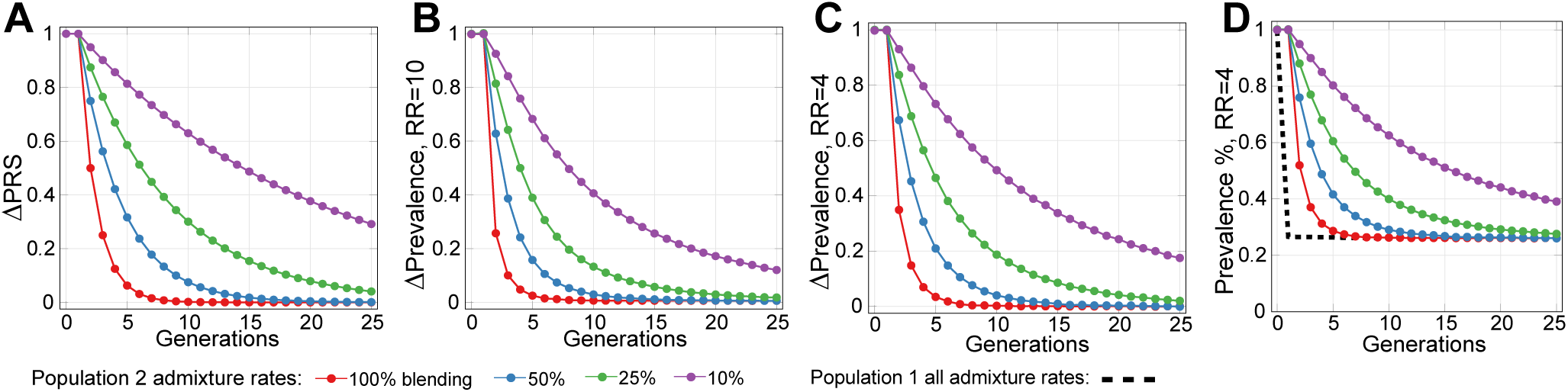
Relative PRS and prevalence progression during population admixture. The normalized relative change in the population PRS and disease prevalence, where “1” is the initial value of the variable, and “0” is the equilibrium value. **(A)** shows the ΔPRS relative to the equilibrium value of the normalized PRS. The displayed fractions are identical to the simple population admixture proportions that would occur at the displayed rates of admixture. **(B)** shows the ΔPrevalence relative to equilibrium normalized disease prevalence progression for population not directly participating in the preventive therapy, when participating population average PRS is maintained at improved tenfold PRS level (RR=10.0, PRS=-2.30). **(C)** shows the ΔPrevalence relative to equilibrium normalized disease prevalence progression for population not directly participating in the preventive therapy, when participating population average PRS is maintained at improved fourfold PRS level (RR=4.0, PRS=-1.386). The normalized relative improvement for population not directly participating in therapy was slightly slower if compared to (B). For 10% admixture rate, the improvement in the first three admixed generations was to the level of 93%, 86% and 80% for RR=4.0 scenario, compared with 92.5%, 84% and 76% for tenfold improvement scenario (RR=10.0) in (B). The higher admixture ratios show even faster prevalence reduction. This shows the comparable relative prevalence reduction, even though the absolute asymptotic reduction differs 2.5 times between these scenarios. **(D)** shows, for comparison with Figure 5D, the absolute prevalence improvement thanks to admixture for populations not participating in preventive gene therapy, when participating population average PRS is maintained at improved fourfold PRS level (RR=4.0, PRS=-1.386); the corresponding normalized figure is depicted in (C).

**Figure A6.**
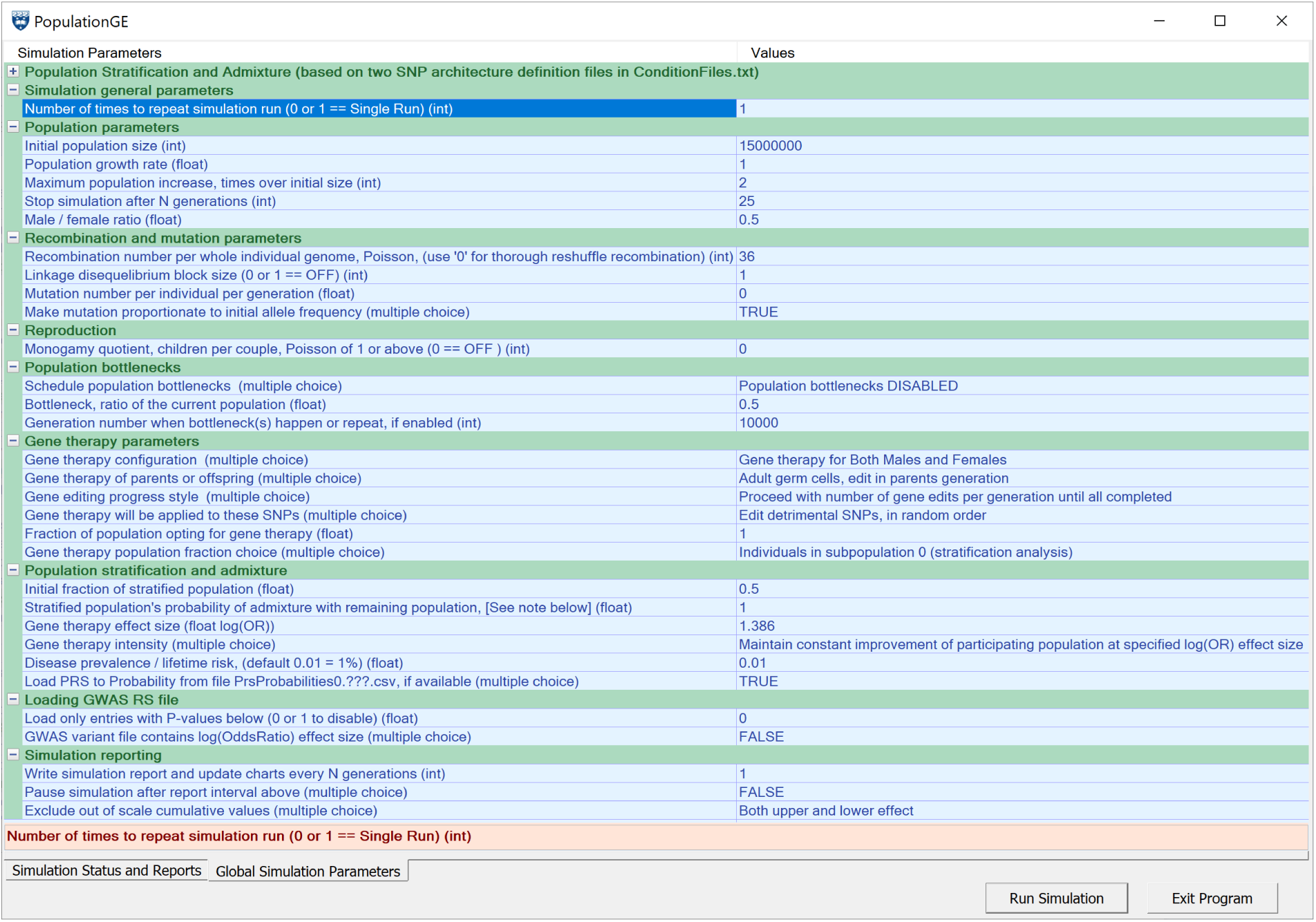
The configuration screen of the simulation program.

## Supplementary Materials

Supplementary Data: A zip file **SupplementaryData**.**ZIP** containing the simulation executable, the source code, R scripts, batch files, and simulation results.

## References

1. Watson, J.D.; Crick, F.H., Molecular Structure of Nucleic Acids: A Structure for Deoxyribose Nucleic Acid. Nature 1953, 171, 737.

2. Morton, N.E.; Crow, J.F.; Muller, H.J., An estimate of the mutational damage in man from data on consanguineous marriages. Proceedings of the National Academy of Sciences 1956, 42, 855–863.

3. Stein, L.D., Human genome: end of the beginning. Nature 2004, 431, 915.

4. Visscher, P.M.; Wray, N.R.; Zhang, Q.; Sklar, P.; McCarthy, M.I.; Brown, M.A.; Yang, J., 10 years of GWAS discovery: biology, function, and translation. The American Journal of Human Genetics 2017, 101, 5–22.

5. OMIM. Available at http://omim.org/statistics/geneMap (accessed June 2, 2019), 2019. URL.

6. Beckmann, J.S.; Estivill, X.; Antonarakis, S.E., Copy number variants and genetic traits: closer to the resolution of phenotypic to genotypic variability. Nature Reviews Genetics 2007, 8, 639.

7. Maroilley, T.; Tarailo-Graovac, M., Uncovering Missing Heritability in Rare Diseases. Genes 2019, 10, 275.

8. Stenson, P.D.; Mort, M.; Ball, E.V.; Evans, K.; Hayden, M.; Heywood, S.; Hussain, M.; Phillips, A.D.; Cooper, D.N., The Human Gene Mutation Database: towards a comprehensive repository of inherited mutation data for medical research, genetic diagnosis and next-generation sequencing studies. Human Genetics 2017, 136, 665–677. doi:10.1007/s00439-017-1779-6.

9. Gao, Z.; Waggoner, D.; Stephens, M.; Ober, C.; Przeworski, M., An estimate of the average number of recessive lethal mutations carried by humans. Genetics 2015, 199, 1243–1254.

10. Chong, J.X.; Buckingham, K.J.; Jhangiani, S.N.; Boehm, C.; Sobreira, N.; Smith, J.D.; Harrell, T.M.; McMillin, M.J.; Wiszniewski, W.; Gambin, T.; others. The genetic basis of Mendelian phenotypes: discoveries, challenges, and opportunities. The American Journal of Human Genetics 2015, 97, 199–215.

11. Ginn, S.L.; Amaya, A.K.; Alexander, I.E.; Edelstein, M.; Abedi, M.R., Gene therapy clinical trials worldwide to 2017: An update. The journal of gene medicine 2018, 20, e3015.

12. Philippidis, A., 25 Up-and-Coming Gene Therapies of 2019. Available at https://www.genengnews.com/a-lists/25-up-and-coming-gene-therapies-of-2019 (accessed June 27, 2019). Genetic Engineering and Biotechnology News (GEN), New York, USA, 2019.

13. Gyngell, C.; Bowman-Smart, H.; Savulescu, J., Moral reasons to edit the human genome: picking up from the Nuffield report. Journal of Medical Ethics 2019. doi:10.1136/medethics-2018-105084.

14. Kemper, J.M.; Gyngell, C.; Savulescu, J., Subsidizing PGD: The Moral Case for Funding Genetic Selection. Journal of Bioethical Inquiry 2019. doi:10.1007/s11673-019-09932-2.

15. Nuffield Council on Bioethics. Genome Editing and Human Reproduction: social and ethical issues; London: Nuffield Council on Bioethics, 2018.

16. Kofler, N.; Kraschel, K.L., Treatment of heritable diseases using CRISPR: Hopes, fears, and reality. Seminars in perinatology. Elsevier, 2018.

17. Pawitan, Y.; Seng, K.C.; Magnusson, P.K., How many genetic variants remain to be discovered? PloS one 2009, 4, e7969.

18. Eyre-Walker, A., Genetic architecture of a complex trait and its implications for fitness and genome-wide association studies. Proceedings of the National Academy of Sciences 2010, 107, 1752–1756.

19. Yang, J.; Ferreira, T.; Morris, A.P.; Medland, S.E.; Madden, P.A.; Heath, A.C.; Martin, N.G.; Montgomery, G.W.; Weedon, M.N.; Loos, R.J., Conditional and joint multiple-SNP analysis of GWAS summary statistics identifies additional variants influencing complex traits. Nature genetics 2012, 44, 369–375.

20. Gonzaga-Jauregui, C.; Lupski, J.R.; Gibbs, R.A., Human genome sequencing in health and disease. Annual review of medicine 2012, 63, 35–61.

21. Lakatta, E.G., So! What’s aging? Is cardiovascular aging a disease? Journal of molecular and cellular cardiology 2015, 83, 1–13.

22. Fuchsberger, C.; Flannick, J.; Teslovich, T.M.; Mahajan, A.; Agarwala, V.; Gaulton, K.J.; Ma, C.; Fontanillas, P.; Moutsianas, L.; McCarthy, D.J.; others. The genetic architecture of type 2 diabetes. Nature 2016, 536, 41–47.

23. Mucci, L.A.; Hjelmborg, J.B.; Harris, J.R.; Czene, K.; Havelick, D.J.; Scheike, T.; Graff, R.E.; Holst, K.; Möller, S.; Unger, R.H.; others. Familial risk and heritability of cancer among twins in Nordic countries. Jama 2016, 315, 68–76.

24. Alzheimer’s Association. 2017 Alzheimer’s disease facts and figures. Alzheimer’s & Dementia 2017, 13, 325–373.

25. Graff, R.E.; Möller, S.; Passarelli, M.N.; Witte, J.S.; Skytthe, A.; Christensen, K.; Tan, Q.; Adami, H.O.; Czene, K.; Harris, J.R., Familial risk and heritability of colorectal cancer in the nordic twin study of cancer. Clinical Gastroenterology and Hepatology 2017, 15, 1256–1264.

26. Fedarko, N.S., Theories and Mechanisms of Aging. In Geriatric Anesthesiology; Nature Publishing Group, 2018; pp. 19–25.

27. Franceschi, C.; Garagnani, P.G.; Morsiani, C.; Conte, M.; Santoro, A.; Grignolio, A.; Monti, D.; Capri, M.; Salvioli, S., The continuum of aging and age-related diseases: common mechanisms but different rates. Frontiers in Medicine 2018, 5, 61.

28. Mitchell, T.J.; Turajlic, S.; Rowan, A.; Nicol, D.; Farmery, J.H.; O’Brien, T.; Martincorena, I.; Tarpey, P.; Angelopoulos, N.; Yates, L.R., Timing the landmark events in the evolution of clear cell renal cell cancer: TRACERx renal. Cell 2018, 173, 611–623.

29. Anderson, C.A.; Soranzo, N.; Zeggini, E.; Barrett, J.C., Synthetic associations are unlikely to account for many common disease genome-wide association signals. PLoS biology 2011, 9, e1000580.

30. Yang, J.; Bakshi, A.; Zhu, Z.; Hemani, G.; Vinkhuyzen, A.A.; Lee, S.H.; Robinson, M.R.; Perry, J.R.; Nolte, I.M.; van Vliet-Ostaptchouk, J.V., Genetic variance estimation with imputed variants finds negligible missing heritability for human height and body mass index. Nature genetics 2015, 47, 1114.

31. Chatterjee, N.; Shi, J.; Garcöa-Closas, M., Developing and evaluating polygenic risk prediction models for stratified disease prevention. Nature Reviews Genetics 2016, 17, 392.

32. Prohaska, A.; Racimo, F.; Schork, A.J.; Sikora, M.; Stern, A.J.; Ilardo, M.; Allentoft, M.E.; Folkersen, L.; Buil, A.; Moreno-Mayar, J.V.; Korneliussen, T.; Geschwind, D.; Ingason, A.; Werge, T.; Nielsen, R.; Willerslev, E., Human Disease Variation in the Light of Population Genomics. Cell 2019, 177, 115–131.

33. Wong, K.H.; Levy-Sakin, M.; Kwok, P.Y., De novo human genome assemblies reveal spectrum of alternative haplotypes in diverse populations. Nature communications 2018, 9, 3040. doi:10.1038/s41467-018-05513-w.

34. Sirugo, G.; Williams, S.M.; Tishkoff, S.A., The Missing Diversity in Human Genetic Studies. Cell 2019, 177, 26–31.

35. Ballouz, S.; Dobin, A.; Gillis, J.A., Is it time to change the reference genome? Genome Biology 2019, 20, 159. doi:10.1186/s13059-019-1774-4.

36. Seyerle, A.A.; Young, A.M.; Jeff, J.M.; Melton, P.E.; Jorgensen, N.W.; Lin, Y.; Carty, C.L.; Deelman, E.; Heckbert, S.R.; Hindorff, L.A.; others. Evidence of heterogeneity by race/ethnicity in genetic determinants of QT interval. Epidemiology (Cambridge, Mass.) 2014, 25, 790.

37. Marigorta, U.M.; Navarro, A., High Trans-ethnic Replicability of GWAS Results Implies Common Causal Variants. PLoS Genetics 2013, 9, e1003566. doi:10.1371/journal.pgen.1003566.

38. Grimsby, J.L.; Porneala, B.C.; Vassy, J.L.; Yang, Q.; Florez, J.C.; Dupuis, J.; Liu, T.; Yesupriya, A.; Chang, M.H.; Ned, R.M.; Dowling, N.F.; Khoury, M.J.; Meigs, J.B.; the MAGIC Investigators. Race-ethnic differences in the association of genetic loci with HbA 1c levels and mortality in US adults: the third National Health and Nutrition Examination Survey (NHANES III). BMC Medical Genetics 2012, 13, 30. doi:10.1186/1471-2350-13-30.

39. Mersha, T.B.; Abebe, T., Self-reported race/ethnicity in the age of genomic research: its potential impact on understanding health disparities. Human Genomics 2015, 9, 1.

40. Belbin, G.M.; Nieves-Colón, M.A.; Kenny, E.E.; Moreno-Estrada, A.; Gignoux, C.R., Genetic diversity in populations across Latin America: implications for population and medical genetic studies. Current opinion in genetics & development 2018, 53, 98–104.

41. Ntzani, E.E.; Liberopoulos, G.; Manolio, T.A.; Ioannidis, J.P., Consistency of genome-wide associations across major ancestral groups. Human genetics 2012, 131, 1057–1071.

42. Martin, A.R.; Gignoux, C.R.; Walters, R.K.; Wojcik, G.L.; Neale, B.M.; Gravel, S.; Daly, M.J.; Bustamante, C.D.; Kenny, E.E., Human Demographic History Impacts Genetic Risk Prediction across Diverse Populations. The American Journal of Human Genetics 2017, 100, 635 – 649. doi:https://doi.org/10.1016/j.ajhg.2017.03.004.

43. Lappalainen, T.; Scott, A.J.; Brandt, M.; Hall, I.M., Genomic Analysis in the Age of Human Genome Sequencing. Cell 2019, 177, 70–84. doi:10.1016/j.cell.2019.02.032.

44. Abel, H.J.; Larson, D.E.; Chiang, C.; Das, I.; Kanchi, K.L.; Layer, R.M.; Neale, B.M.; Salerno, W.J.; Reeves, C.; Buyske, S.; NHGRI Centers for Common Disease Genomics.; Matise, T.C.; Muzny, D.M.; Zody, M.C.; Lander, E.S.; Dutcher, S.K.; Stitziel, N.O.; Hall, I.M., Mapping and characterization of structural variation in 17,795 deeply sequenced human genomes. bioRxiv 2018. doi:10.1101/508515.

45. Zook, J.M.; Hansen, N.F.; Olson, N.D.; Chapman, L.M.; Mullikin, J.C.; Xiao, C.; Sherry, S.; Koren, S.; Phillippy, A.M.; Boutros, P.C.; Sahraeian, S.M.E.; Huang, V.; Rouette, A.; Alexander, N.; Mason, C.E.; Hajirasouliha, I.; Ricketts, C.; Lee, J.; Tearle, R.; Fiddes, I.T.; Barrio, A.M.; Wala, J.; Carroll, A.; Ghaffari, N.; Rodriguez, O.L.; Bashir, A.; Jackman, S.; Farrell, J.J.; Wenger, A.M.; Alkan, C.; Soylev, A.; Schatz, M.C.; Garg, S.; Church, G.; Marschall, T.; Chen, K.; Fan, X.; English, A.C.; Rosenfeld, J.A.; Zhou, W.; Mills, R.E.; Sage, J.M.; Davis, J.R.; Kaiser, M.D.; Oliver, J.S.; Catalano, A.P.; Chaisson, M.J.; Spies, N.; Sedlazeck, F.J.; Salit, M.; The Genome in a Bottle Consortium. A robust benchmark for germline structural variant detection. bioRxiv 2019. doi:10.1101/664623.

46. Oliynyk, R.T., Quantifying the Potential for Future Gene Therapy to Lower Lifetime Risk of Polygenic Late-Onset Diseases. International Journal of Molecular Sciences 2019, 20. doi:10.3390/ijms20133352.

47. Falconer, D.S., The inheritance of liability to certain diseases, estimated from the incidence among relatives. Annals of human genetics 1965, 29, 51–76.

48. Falconer, D., The inheritance of liability to diseases with variable age of onset, with particular reference to diabetes mellitus. Annals of human genetics 1967, 31, 1–20.

49. Wray, N.R.; Goddard, M.E., Multi-locus models of genetic risk of disease. Genome Medicine 2010, 2, 10.

50. Oliynyk, R.T., Age-related late-onset disease heritability patterns and implications for genome-wide association studies. PeerJ 2019, 7, e7168. doi:10.7717/peerj.7168.

51. Cox, D., Regression Models and Life-Tables. Journal of the Royal Statistical Society. Series B (Methodological) 1972, 34, 187–220.

52. Mars, N.J.; Koskela, J.T.; Ripatti, P.; Kiiskinen, T.T.; Havulinna, A.S.; Lindbohm, J.V.; Ahola-Olli, A.; Kurki, M.; Karjalainen, J.; Palta, P.; FinnGen.; Neale, B.M.; Daly, M.; Salomaa, V.; Palotie, A.; Widen, E.; Ripatti, S., Polygenic and clinical risk scores and their impact on age at onset of cardiometabolic diseases and common cancers. bioRxiv 2019. doi:10.1101/727057.

53. Vicente, C.T.; Revez, J.A.; Ferreira, M.A.,R. Lessons from ten years of genome-wide association studies of asthma. Clinical & Translational Immunology 2017, 6, e165. doi:10.1038/cti.2017.54.

54. Willis-Owen, S.A.; Cookson, W.O.; Moffatt, M.F., The Genetics and Genomics of Asthma. Annual Review of Genomics and Human Genetics 2018, 19, 223–246. PMID: 30169121, doi:10.1146/annurev-genom-083117-021651.

55. Lipton, R.B.; Bigal, M.E.; Diamond, M.; Freitag, F.; Reed, M.; Stewart, W.F.; others. Migraine prevalence, disease burden, and the need for preventive therapy. Neurology 2007, 68, 343–349.

56. Chalmer, M.A.; Esserlind, A.L.; Olesen, J.; Hansen, T.F., Polygenic risk score: use in migraine research. The journal of headache and pain 2018, 19, 29–29. doi:10.1186/s10194-018-0856-0.

57. Riesmeijer, S.A.; Werker, P.M.; Nolte, I.M., Ethnic differences in prevalence of Dupuytren disease can partly be explained by known genetic risk variants. European Journal of Human Genetics 2019, p. 1.

58. Kurkó, J.; Besenyei, T.; Laki, J.; Glant, T.T.; Mikecz, K.; Szekanecz, Z., Genetics of rheumatoid arthritis - a comprehensive review. Clinical reviews in allergy & immunology 2013, 45, 170–179.

59. Kuo, C.F.; Grainge, M.J.; Valdes, A.M.; See, L.C.; Luo, S.F.; Yu, K.H.; Zhang, W.; Doherty, M., Familial aggregation of systemic lupus erythematosus and coaggregation of autoimmune diseases in affected families. JAMA internal medicine 2015, 175, 1518–1526.

60. of the Psychiatric Genomics Consortium, S.W.G.; others. Genomic dissection of bipolar disorder and schizophrenia, including 28 subphenotypes. Cell 2018, 173, 1705–1715.

61. Liu, J.Z.; Anderson, C.A., Genetic studies of Crohn’s disease: past, present and future. Best Practice & Research Clinical Gastroenterology 2014, 28, 373–386.

62. Lee, J.C.; Biasci, D.; Roberts, R.; Gearry, R.B.; Mansfield, J.C.; Ahmad, T.; Prescott, N.J.; Satsangi, J.; Wilson, D.C.; Jostins, L.; Anderson, C.A.; Traherne, J.A.; Lyons, P.A.; Parkes, M.; Smith, K.G.C. Genome-wide association study identifies distinct genetic contributions to prognosis and susceptibility in Crohn’s disease. Nature genetics 2017, 49, 262–269B.

63. Gluckman, P.D.; Low, F.M.; Buklijas, T.; Hanson, M.A.; Beedle, A.S., How evolutionary principles improve the understanding of human health and disease. Evolutionary Applications 2011, 4, 249–263.

64. Bergen, S.E.; O’Dushlaine, C.T.; Lee, P.H.; Fanous, A.H.; Ruderfer, D.M.; Ripke, S.; International Schizophrenia Consortium, S.S.C.; Sullivan, P.F.; Smoller, J.W.; Purcell, S.M.; Corvin, A., Genetic modifiers and subtypes in schizophrenia: investigations of age at onset, severity, sex and family history. Schizophrenia research 2014, 154, 48–53. doi:10.1016/j.schres.2014.01.030.

65. Wang, K.; Gaitsch, H.; Poon, H.; Cox, N.J.; Rzhetsky, A., Classification of common human diseases derived from shared genetic and environmental determinants. Nature genetics 2017, 49, 1319–1325. doi:10.1038/ng.3931.

66. Polubriaginof, F.C.; Vanguri, R.; Quinnies, K.; Belbin, G.M.; Yahi, A.; Salmasian, H.; Lorberbaum, T.; Nwankwo, V.; Li, L.; Shervey, M.M.; Glowe, P.; Ionita-Laza, I.; Simmerling, M.; Hripcsak, G.; Bakken, S.; Goldstein, D.; Kiryluk, K.; Kenny, E.E.; Dudley, J.; Vawdrey, D.K.; Tatonetti, N.P., Disease heritability inferred from familial relationships reported in medical records. Cell 2018, 173, 1692–1704.

67. Gravel, S., Population genetics models of local ancestry. Genetics 2012, 191, 607–619.

68. Huber, M.; Chen, Y.; Dinwoodie, I.; Dobra, A.; Nicholas, M., Monte carlo algorithms for Hardy–Weinberg proportions. Biometrics 2006, 62, 49–53.

69. Mayo, O., A century of Hardy–Weinberg equilibrium. Twin Research and Human Genetics 2008, 11, 249–256.

70. Chakraborty, R.; Weiss, K.M., Frequencies of complex diseases in hybrid populations. American journal of physical anthropology 1986, 70, 489–503.

71. Borzecki, A.M.; Bridgers, D.K.; Liebschutz, J.M.; Kader, B.; Kazis, S.L.E.; Berlowitz, D.R., Racial differences in the prevalence of atrial fibrillation among males. Journal of the National Medical Association 2008, 100, 237–246.

72. Larsen, S.; Krogsgaard, D.; Larsen, L.A.; Iachina, M.; Skytthe, A.; Frederiksen, H., Genetic and environmental influences in Dupuytren’s disease: a study of 30,330 Danish twin pairs. Journal of Hand Surgery (European Volume) 2015, 40, 171–176.

73. Lee, K.H.; Kim, J.H.; Lee, C.H.; Kim, S.J.; Jo, Y.H.; Lee, M.; Choi, W.S., The epidemiology of Dupuytren’s disease in Korea: a nationwide population-based study. Journal of Korean medical science 2018, 33.

74. Yeh, C.C.; Huang, K.F.; Ho, C.H.; Chen, K.T.; Liu, C.; Wang, J.J.; Chu, C.C., Epidemiological profile of Dupuytren’s disease in Taiwan (Ethnic Chinese): a nationwide population-based study. BMC musculoskeletal disorders 2015, 16, 20.

75. Molokhia, M.; McKeigue, P., Risk for rheumatic disease in relation to ethnicity and admixture. Arthritis Research & Therapy 2000, 2, 115.

76. Chen, L.; Morris, D.L.; Vyse, T.J., Genetic advances in systemic lupus erythematosus: an update. Current opinion in rheumatology 2017, 29, 423–433.

77. Lim, S.S.; Bayakly, A.R.; Helmick, C.G.; Gordon, C.; Easley, K.A.; Drenkard, C., The incidence and prevalence of systemic lupus erythematosus, 2002–2004: the Georgia Lupus Registry. Arthritis & rheumatology 2014, 66, 357–368.

78. Herráez, D.L.; Martönez-Bueno, M.; Riba, L.; de la Torre, I.G.; Sacnún, M.; Goñi, M.; Berbotto, G.A.; Paira, S.; Musuruana, J.L.; Graf, C.E.; others. Rheumatoid arthritis in Latin Americans enriched for Amerindian ancestry is associated with loci in chromosomes 1, 12, and 13, and the HLA class II region. Arthritis & Rheumatism 2013, 65, 1457–1467.

79. Dudbridge, F., Polygenic epidemiology. Genetic epidemiology 2016, 40, 268–272.

80. Hormozdiari, F.; Zhu, A.; Kichaev, G.; Ju, C.J.T.; Segrè, A.V.; Joo, J.W.J.; Won, H.; Sankararaman, S.; Pasaniuc, B.; Shifman, S.; others. Widespread allelic heterogeneity in complex traits. The American Journal of Human Genetics 2017, 100, 789–802. doi:10.1016/j.ajhg.2017.04.005.

81. Salzano, F.M.; Sans, M., Interethnic admixture and the evolution of Latin American populations. Genetics and molecular biology 2014, 37, 151–170.

82. Acuna-Hidalgo, R.; Veltman, J.A.; Hoischen, A., New insights into the generation and role of de novo mutations in health and disease. Genome biology 2016, 17, 241.

83. Lynch, M., Mutation and human exceptionalism: our future genetic load. Genetics 2016, 202, 869–875.

84. Lynch, M.; Ackerman, M.S.; Gout, J.F.; Long, H.; Sung, W.; Thomas, W.K.; Foster, P.L., Genetic drift, selection and the evolution of the mutation rate. Nature Reviews Genetics 2016, 17, 704–714.

85. Gao, Z.; Wyman, M.J.; Sella, G.; Przeworski, M., Interpreting the dependence of mutation rates on age and time. PLoS biology 2016, 14, e1002355.

86. Engels, W.R., Exact tests for Hardy–Weinberg proportions. Genetics 2009, 183, 1431–1441.

87. Shifman, S.; Kuypers, J.; Kokoris, M.; Yakir, B.; Darvasi, A., Linkage disequilibrium patterns of the human genome across populations. Human molecular genetics 2003, 12, 771–776.

88. Martin, E.R.; Tunc, I.; Liu, Z.; Slifer, S.H.; Beecham, A.H.; Beecham, G.W., Properties of global-and local-ancestry adjustments in genetic association tests in admixed populations. Genetic epidemiology 2018, 42, 214–229.

89. Risch, N.; Merikangas, K., The future of genetic studies of complex human diseases. Science 1996, 273, 1516–1517.

90. Lu, Q.; Elston, R.C., Using the optimal receiver operating characteristic curve to design a predictive genetic test, exemplified with type 2 diabetes. The American Journal of Human Genetics 2008, 82, 641–651.

91. Stearns, F.W., One hundred years of pleiotropy: a retrospective. Genetics 2010, 186, 767–773.

92. Paaby, A.B.; Rockman, M.V., The many faces of pleiotropy. Trends in Genetics 2013, 29, 66–73.

93. Bhatia, G.; Patterson, N.; Sankararaman, S.; Price, A.L., Estimating and interpreting FST: the impact of rare variants. Genome research 2013, 23, 1514–1521.

94. Berner, D., Allele Frequency Difference AFD-An Intuitive Alternative to FST for Quantifying Genetic Population Differentiation. Genes 2019, 10. doi:10.3390/genes10040308.

95. Wray, N.R.; Goddard, M.E.; Visscher, P.M., Prediction of individual genetic risk to disease from genome-wide association studies. Genome research 2007, 17, 1520–1528.

96. Martin, A.R.; Daly, M.J.; Robinson, E.B.; Hyman, S.E.; Neale, B.M., Predicting polygenic risk of psychiatric disorders. Biological psychiatry 2018.

97. Gejman, P.V.; Sanders, A.R.; Duan, J., The role of genetics in the etiology of schizophrenia. The Psychiatric clinics of North America 2010, 33, 35–66. doi:10.1016/j.psc.2009.12.003.

98. Ferreira, M.A.; Mathur, R.; Vonk, J.M.; Szwajda, A.; Brumpton, B.; Granell, R.; Brew, B.K.; Ullemar, V.; Lu, Y.; Jiang, Y.; Magnusson, P.K.; Karlsson, R.; Hinds, D.A.; Paternoster, L.; Koppelman, G.H.; Almqvist, C., Genetic Architectures of Childhood-and Adult-Onset Asthma Are Partly Distinct. The American Journal of Human Genetics 2019, 104, 665–684. doi:https://doi.org/10.1016/j.ajhg.2019.02.022.

99. Pardiñas, A.F.; Holmans, P.; Pocklington, A.J.; Escott-Price, V.; Ripke, S.; Carrera, N.; Legge, S.E.; Bishop, S.; Cameron, D.; Hamshere, M.L.; others. Common schizophrenia alleles are enriched in mutation-intolerant genes and in regions under strong background selection. Nature genetics 2018, 50, 381. doi:10.1038/s41588-018-0059-2.

100. Wray, N.R.; Visscher, P.M., Narrowing the boundaries of the genetic architecture of schizophrenia. Schizophrenia bulletin 2009, 36, 14–23. doi:10.1093/schbul/sbp137.

101. Ugowe, F.E.; Jackson, L.R.I.; Thomas, K.L., Racial and ethnic differences in the prevalence, management, and outcomes in patients with atrial fibrillation: A systematic review. Heart Rhythm 2018, 15, 1337–1345. doi:10.1016/j.hrthm.2018.05.019.

102. Musunuru, K.; Kathiresan, S., Genetics of Common, Complex Coronary Artery Disease. Cell 2019, 177, 132–145.

103. Stern, M.C.; Fejerman, L.; Das, R.; Setiawan, V.W.; Cruz-Correa, M.R.; Perez-Stable, E.J.; Figueiredo, J.C., Variability in Cancer Risk and Outcomes Within US Latinos by National Origin and Genetic Ancestry. Current Epidemiology Reports 2016, 3, 181–190. doi:10.1007/s40471-016-0083-7.

104. Holley, A.; Northcott, H.; Gladding, P.; Harding, S.; Larsen, P., Significant Differences in Genetic Risk Profiles Between Maori and European Presenting with Myocardial Infarction. Heart, Lung and Circulation 2017, 26, S307–S308.

105. Gurdasani, D.; Carstensen, T.; Tekola-Ayele, F.; Pagani, L.; Tachmazidou, I.; Hatzikotoulas, K.; Karthikeyan, S.; Iles, L.; Pollard, M.O.; Choudhury, A.; Ritchie, G.R.S.; Xue, Y.; Asimit, J.; Nsubuga, R.N.; Young, E.H.; Pomilla, C.; Kivinen, K.; Rockett, K.; Kamali, A.; Doumatey, A.P.; Asiki, G.; Seeley, J.; Sisay-Joof, F.; Jallow, M.; Tollman, S.; Mekonnen, E.; Ekong, R.; Oljira, T.; Bradman, N.; Bojang, K.; Ramsay, M.; Adeyemo, A.; Bekele, E.; Motala, A.; Norris, S.A.; Pirie, F.; Kaleebu, P.; Kwiatkowski, D.; Tyler-Smith, C.; Rotimi, C.; Zeggini, E.; Sandhu, M.S., The African Genome Variation Project shapes medical genetics in Africa. Nature 2014, 517, 327 EP –.

106. Zanetti, D.; Weale, M.E., Transethnic differences in GWAS signals: A simulation study. Annals of Human Genetics 2018, 82, 280–286. doi:10.1111/ahg.12251.

107. National Academies of Sciences, Engineering, and Medicine. Human genome editing: Science, ethics, and governance; National Academies Press, 2017.

108. Maurano, M.T.; Humbert, R.; Rynes, E.; Thurman, R.E.; Haugen, E.; Wang, H.; Reynolds, A.P.; Sandstrom, R.; Qu, H.; Brody, J.; Shafer, A.; Neri, F.; Lee, K.; Kutyavin, T.; Stehling-Sun, S.; Johnson, A.K.; Canfield, T.K.; Giste, E.; Diegel, M.; Bates, D.; Hansen, R.S.; Neph, S.; Sabo, P.J.; Heimfeld, S.; Raubitschek, A.; Ziegler, S.; Cotsapas, C.; Sotoodehnia, N.; Glass, I.; Sunyaev, S.R.; Kaul, R.; Stamatoyannopoulos, J.A., Systematic Localization of Common Disease-Associated Variation in Regulatory DNA. Science 2012, 337, 1190–1195. doi:10.1126/science.1222794.

109. Lee, D.; Gorkin, D.U.; Baker, M.; Strober, B.J.; Asoni, A.L.; McCallion, A.S.; Beer, M.A., A method to predict the impact of regulatory variants from DNA sequence. Nature genetics 2015, 47, 955.

110. Zhu, Y.; Tazearslan, C.; Suh, Y., Challenges and progress in interpretation of non-coding genetic variants associated with human disease. Experimental Biology and Medicine 2017, 242, 1325–1334.

111. Dong, C.; Wei, P.; Jian, X.; Gibbs, R.; Boerwinkle, E.; Wang, K.; Liu, X., Comparison and integration of deleteriousness prediction methods for nonsynonymous SNVs in whole exome sequencing studies. Human molecular genetics 2014, 24, 2125–2137.

112. Ioannidis, N.M.; Rothstein, J.H.; Pejaver, V.; Middha, S.; McDonnell, S.K.; Baheti, S.; Musolf, A.; Li, Q.; Holzinger, E.; Karyadi, D.; others. REVEL: an ensemble method for predicting the pathogenicity of rare missense variants. The American Journal of Human Genetics 2016, 99, 877–885.

113. Jian, X.; Liu, X., In Silico Prediction of Deleteriousness for Nonsynonymous and Splice-Altering Single Nucleotide Variants in the Human Genome. In In Vitro Mutagenesis; Springer, 2017; pp. 191–197.

114. Wagih, O.; Galardini, M.; Busby, B.P.; Memon, D.; Typas, A.; Beltrao, P., A resource of variant effect predictions of single nucleotide variants in model organisms. Molecular systems biology 2018, 14, e8430.

115. Yauy, K.; Baux, D.; Pegeot, H.; Van Goethem, C.; Mathieu, C.; Guignard, T.; Morales, R.J.; Lacourt, D.; Krahn, M.; Lehtokari, V.L.; others. MoBiDiC Prioritization Algorithm, a Free, Accessible, and Efficient Pipeline for Single-Nucleotide Variant Annotation and Prioritization for Next-Generation Sequencing Routine Molecular Diagnosis. The Journal of Molecular Diagnostics 2018, 20, 465–473.

116. Korvigo, I.; Afanasyev, A.; Romashchenko, N.; Skoblov, M., Generalising better: Applying deep learning to integrate deleteriousness prediction scores for whole-exome SNV studies. PloS one 2018, 13, e0192829.

117. Wright, A.V.; Nuñez, J.K.; Doudna, J.A., Biology and applications of CRISPR systems: harnessing nature’s toolbox for genome engineering. Cell 2016, 164, 29–44.

118. Carroll, D., Genome engineering with zinc-finger nucleases. Genetics 2011, 188, 773–782.

119. Joung, J.K.; Sander, J.D., TALENs: a widely applicable technology for targeted genome editing. Nature reviews Molecular cell biology 2013, 14, 49.

120. Kocak, D.D.; Josephs, E.A.; Bhandarkar, V.; Adkar, S.S.; Kwon, J.B.; Gersbach, C.A., Increasing the specificity of CRISPR systems with engineered RNA secondary structures. Nature biotechnology 2019, 37, 657.

121. Smith, C.J.; Castanon, O.; Said, K.; Volf, V.; Khoshakhlagh, P.; Hornick, A.; Ferreira, R.; Wu, C.T.; Güell, M.; Garg, S.; Myllykallio, H.; Church, G.M., Enabling large-scale genome editing by reducing DNA nicking. bioRxiv 2019. doi:10.1101/574020.

122. Strecker, J.; Ladha, A.; Gardner, Z.; Schmid-Burgk, J.L.; Makarova, K.S.; Koonin, E.V.; Zhang, F., RNA-guided DNA insertion with CRISPR-associated transposases. Science 2019. doi:10.1126/science.aax9181.

123. Thompson, D.; Aboulhouda, S.; Hysolli, E.; Smith, C.; Wang, S.; Castanon, O.; Church, G., The future of multiplexed eukaryotic genome engineering. ACS chemical biology 2017, 13, 313–325.

124. Kohman, R.E.; Kunjapur, A.M.; Hysolli, E.; Wang, Y.; Church, G.M., From Designing the Molecules of Life to Designing Life: Future Applications Derived from Advances in DNA Technologies. Angewandte Chemie 2018, 57(16), 4313–4328.

125. Noh, M.; Yip, B.; Lee, Y.; Pawitan, Y., Multicomponent variance estimation for binary traits in family-based studies. Genetic epidemiology 2006, 30, 37–47.

